# Evolution of enzyme levels in metabolic pathways: A theoretical approach. Part 2

**DOI:** 10.1101/2021.11.18.469121

**Authors:** Charlotte Coton, Christine Dillmann, Dominique de Vienne

## Abstract

Metabolism is essential for cell function and adaptation. Because of their central role in metabolism, kinetic parameters and enzyme concentrations are under constant selective pressure to adapt the fluxes of the metabolic networks to the needs of the organism. In the line of various studies dealing with enzyme evolution, we recently developed a model of evolution of enzyme concentrations under selection for increased flux, considered as a proxy of fitness (Coton et al. 2021). Taking into account two realistic cellular constraints, competition for resources and co-regulations, we determined the evolutionary equilibria and the ranges of neutral variations of enzyme concentrations. In this article, we give more generality to this model, by considering that the enzymes of a pathway can belong to different groups of co-regulation. We determined the equilibria and showed that the constraints modify the adaptive landscape by limiting the number of independent dimensions. We also showed that any trade-off between enzyme concentration is sufficient to limit the flux and to relax selection for increasing other enzyme concentrations. Even though the model is based on simplifying assumptions, the complexity of the relationship between enzyme concentrations prevents the analysis of selective neutrality.

## 1 Introduction

The key role of metabolism in cell functioning and adaptation accounts for the large number of studies devoted to its evolution. The main lines of research concern the evolution of metabolic network structure and properties (Noda-Garcia et al. 2018; Newton et al. 2018; Sambamoorthy et al. 2019; Morrison and Badyaev 2017), the polymorphism and rate of evolution of enzyme coding genes (Larracuente et al. 2008; Rausher et al. 1999; Colombo et al. 2014; Flowers et al. 2007; Sellis and Longo 2015) and the evolution of enzyme properties (Eanes 1999; Kacser and Beeby 1984; Serohijos and Shakhnovich 2014; Bershtein et al. 2017; Heckmann et al. 2018). In this context, it is striking to observe that very few studies have addressed the question of the evolution of enzyme concentrations in response to selection on the metabolic flux, considered to be a proxy of fitness. Yet, quantitative proteomics in various species has revealed the extent of the genetic variability of enzyme concentrations (Damerval et al. 1994; Blein-Nicolas et al. 2013; Chick et al. 2016; Jiang et al. 2019).

In a classical paper, Hartl et al. (1985) analysed the evolution of enzyme quantity, equated to activity, under selection for higher metabolic flux. Relying on the metabolic control theory (MCT) (Kacser and Burns 1973; Heinrich and Rapoport 1974), they proposed an explanation for the selective neutrality of molecular polymorphism. Under selection, the mutations that increase the flux are progressively fixed, but due to the concavity of the enzyme-flux relationship, the selection coefficients of the mutations become smaller and smaller. At the plateau of the curve, enzyme variations become nearly-neutral.

This interesting model relied on two quite restrictive assumptions: (i) the mutations always affect the same enzyme of the pathway; (ii) there is no limitation to enzyme concentrations, in spite of the well-documented constraints on energy and space in the cell (Kurland and Dong 1996; Ellis 2001; Klumpp et al. 2019). In addition this model has never be tested following an adaptive dynamics approach.

In a companion paper, we recently filled these gaps by developing the first adaptive dynamics model of the evolution of enzyme concentrations in linear pathways (Coton et al. 2021). We considered jointly the evolution of all enzymes, because the concentration of any enzyme of the pathway can be targeted by mutation, and we introduced in the model possible competition for cellular resources and/or co-regulations, which prevents enzyme concentrations from varying freely (Heinrich et al. 1991; Lion et al. 2004).

We wrote a system of differential equations that describes the evolution of enzyme concentrations and we analysed the enzyme properties and constraint-specific parameters that shape the adaptive landscape. We showed that the relative enzyme concentrations reach evolutionary equilibria that depend on the kinetic parameters of enzymes and thermodynamic constants, and which are modified by competition and/or co-regulation. We also analysed the evolution of the flux upon selection and determined its limit value in the case of competition and/or negative co-regulation. The values of the diverse equilibria and of the flux, predicted analytically or numerically, were confirmed by computer simulations of long term evolution. Finally we showed that the selective neutrality of all enzymes is achieved at equilibrium whatever the constraints, but the range of neutral variation of concentrations vary in a large extent, depending on constraint-specific parameters. In the case of co-regulations, initial enzyme concentrations affect the evolutionary outcome.

In the previous developments, we considered as a first analysis that the enzymes of a pathway are either independent or all co-regulated to each other, with or without competition. However, enzyme concentrations of a given pathway can be differently regulated. For example, in *Saccharomyces cerevisiae* many transcription factors (TFs) control the expression of glycolytic genes, in different ways depending on the promoters (Chambers et al. 1995). In the pathogenic yeast *Candida albicans*, two major TFs, Gal4p and Tye7p, activate the genes of the glycolytic pathway: both control diverse enzymes, but upstream enzyme genes are mainly activated by Tye7p (Askew et al. 2009). In *Escherichia coli*, only about 30 % of sequential enzymes in pathways are both regulated, including 75 % (down to 8 % for enzymes at junction) that share the same TF, *i*.*e*. are co-regulated (Seshasayee et al. 2009). Also in *Escherichia coli*, Goelzer et al. (2008) linked the different metabolic pathways to the regulatory networks, and shown that enzymes can be controlled by different TFs (activators or repressors) which can be common to other enzymes, or not regulated at all.

In this work, we propose a generalization of the enzyme evolution model of Coton et al. (2021). We take into account the realistic situation where different groups of coregulated enzymes coexist in a pathway, with or without competition between all enzymes for cellular resources. To search the equilibria we decompose the relative concentrations into intra- and inter-group relative concentrations. We found the evolutionary equilibria when there is no competition, which differ according to the type of co-regulation – positive or negative – within the groups. When there is competition, we did not find the evolutionary equilibria analytically, but the simulations show that they exist. Finally, we determine that mutations become neutral around the equilibrium, but the complexity of the relationships between enzyme concentrations prevent us to compute the range of neutral variations. We show that increasing the number of co-regulations between enzymes diminishes the number of degrees of freedom of the system, which reduces the extent of the adaptive landscape. Thus, only two negatively co-regulated enzymes are sufficient to limit the flux, which results in a progressively decrease of the selective pressure on all other enzymes. Moreover the presence of co-regulations means that the outcome of the evolutionary process depends on the initial concentrations.

## 2 Material & Methods

The modelling approach and simulation methods are developed in Coton et al. (2021). Hereafter we present them briefly and we complement the theoretical background to treat the general case where the enzymes of the pathway belong to different groups of co-regulation. The glossary of mathematical symbols is in appendix A.

### 2.1 Theoretical background

#### 2.1.1 Flux expression

In a linear metabolic pathway of *n* unsaturated Michaelian-Menten enzymes, the flux at the steady state is (Kacser and Burns 1981):

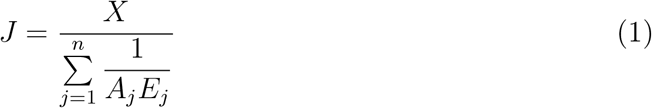

where:

- *X* is a constant.
- *A*_*j*_ is called *pseudo-activity* of enzyme *j. A*_*j*_ is a composite parameter which includes the catalytic constant *k*_cat,*j*_ and the Michaelis-Menten constant 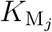 of enzyme *j*, and the product *K*_1,*j*−1_ of the equilibrium constants of the reactions upstream enzyme *j*:

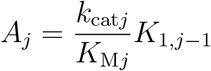
- *E*_*j*_ is the concentration of enzyme *j*.

#### 2.1.2 Formalizing the dependence between enzymes

##### The redistribution coefficients

Constraints on enzyme concentrations, such as co-regulations or competition for resources, create interdependence between enzymes. As a consequence, a mutation that changes the concentration of a target enzyme indirectly changes the concentration of the other enzymes. We model this effect by the following relations. For a target enzyme *i*, we have

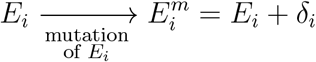

where *δ*_*i*_ is the actual effect of the mutation targeting enzyme *i*. For the other enzymes *j* ≠ *i*, we have

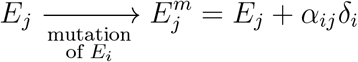

where *α*_*ij*_ is the *redistribution coefficient*, the value of which depends on the type of constraint applied (Lion et al. 2004; Coton et al. 2021):

1. Without any constraint, *α*_*ij*_ is null.
2. In the case of co-regulation only, *α*_*ij*_ is equal to *β*_*ij*_, the *co-regulation coefficient*, which expresses the change in concentration of enzyme *j* due to co-regulation when concentration of enzyme *i* varies. We assumed linearity between co-regulated enzymes, so we posed: *β*_*ii*_ = 1, *β*_*ij*_ = 1*/β*_*ji*_ and, for any triplet of co-regulated enzymes (*i, j, k*), *β*_*ij*_ = *β*_*ik*_*β*_*kj*_. If enzymes *i* and *j* are not co-regulated, we have *β*_*ij*_ = *β*_*ji*_ = 0. All pairwise co-regulation coefficients can be grouped in a coregulation matrix, noted **M**_*β*_. The co-regulation matrix is assumed to result from the action of a hypothetical gene regulatory network. The *global co-regulation coefficient* 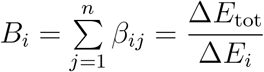, quantifies the impact due to co-regulation of a variation of *E*_*i*_ on the total enzyme concentration *E*_tot_, where 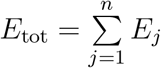.
3. In the case of competition, *E*_tot_ is fixed and 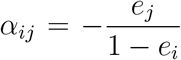, where *e*_*i*_ (resp. *e*_*j*_) is the relative enzyme concentration: 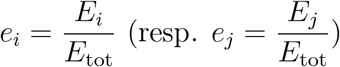.
4. In the case of co-regulation *and* competition, 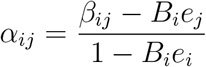.

Note that ∀*i, α*_*ii*_ = 1.

##### The flux response coefficient

When there are dependencies between enzyme concentrations in a pathway, the classical flux control coefficient (Kacser et al. 1995) cannot be used to quantify the effect on the flux of the variation of an enzyme concentration. So we used the flux response coefficient (de Vienne et al. 2001; Coton et al. 2021) that includes not only the parameters of all enzymes, but also the redistribution coefficients *α*_*ij*_:

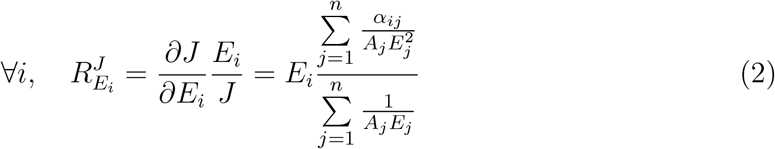

### 2.2 Co-regulation groups

In Coton et al. (2021), we assumed that in the case of co-regulation, *all* enzymes of the pathway are co-regulated, *i*.*e. β*_*ij*_ ≠ 0 for all *i* ≠ *j*. Here we consider the more realistic situation where there are different groups of co-regulated enzymes.

Let Φ_*q*_ the co-regulation group *q*, which includes *m*_*q*_ enzymes. With *p* co-regulation groups, we have 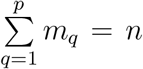, with *n* the total number of enzymes. Within a group the enzymes are co-regulated, and are independent from all enzymes outside the group. This means that for all enzyme *i* ∈ Φ_*q*_, *β*_*ij*_ ≠ 0 if *j* ∈ Φ_*q*_ and *β*_*ij*_ = 0 = *β*_*ji*_ if *j ∉* Φ_*q*_. Thus some *β*_*ij*_ are null in the co-regulation matrix **M**_*β*_.

We defined three possible types of co-regulation groups, denoted by the indicator variable *θ*_*q*_:

i. if there are only positive co-regulations, *θ*_*q*_ = +1 and Φ_*q*_ is called a *positive group*;
ii. if there is at least one negative co-regulation in the group, *θ*_*q*_ =−1 and Φ_*q*_ is called a *negative group*;
iii. if there is only one enzyme in Φ_*q*_, *θ*_*q*_ = 0 and Φ_*q*_ is called a *singleton* (the enzyme is independent from all other enzymes).

Note that the situations analysed by Coton et al. (2021) are particular cases of this general co-regulation model: co-regulation between *all* enzymes corresponds to *p* = 1 and *m*_1_ = *n*, and independence between all enzymes to *p* = *n* and *m*_*q*_ = 1 for all *q*.

Figure 1 shows a ten-enzyme pathway with three co-regulation groups and two singletons and the corresponding co-regulation matrix.

**Figure 1.**
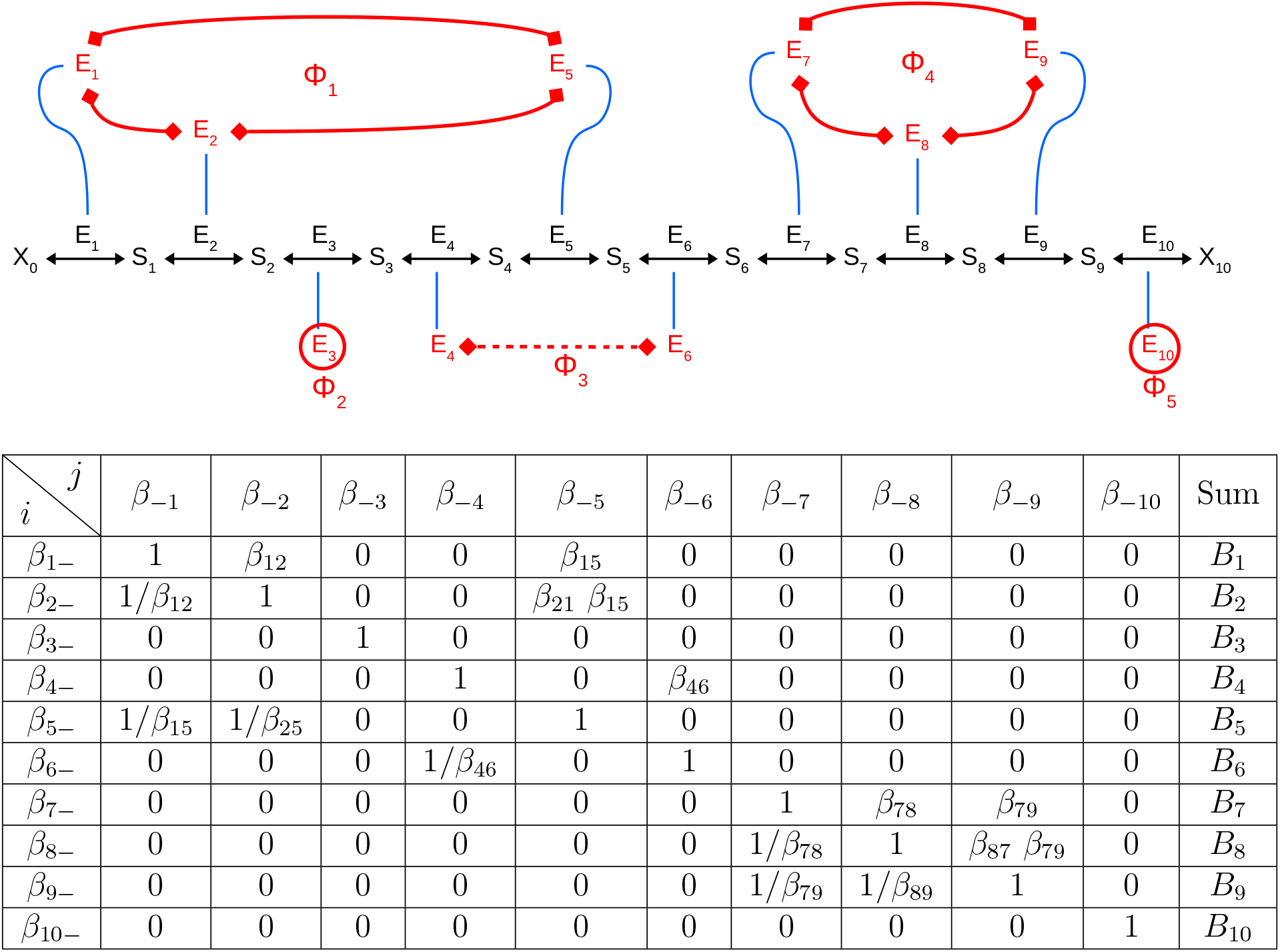
Example of co-regulation groups in a linear pathway. The pathway transforms an initial substrate X_0_ in a final product X_10_ through ten reversible reactions catalysed by enzymes E_1_ to E_10_. S_*i*_ is the product of enzyme E_*i*_ and the substrate of enzyme E_*i*+1_. There are three co-regulation groups and two singletons (*p* = 5), noted Φ_1_ to Φ_5_, which result from hypothetical gene regulatory processes (in red). We have Φ_1_ = {1, 2, 5} ; Φ_2_ = {3} ; Φ_3_ = {4, 6}; Φ_4_ = {7, 8, 9} ; and Φ_5_ = {10}. The blue lines relate genes g_1_ to g_10_ to enzymes E_1_ to E_10_, respectively. Red plain lines and red dashed lines denote positive and negative co-regulations, respectively. The singletons (circled enzymes) are not co-regulated with any other enzyme. The table shows the co-regulation matrix of this pathway. This table includes the sub-matrices for each of the three co-regulation groups: Φ_1_ = {1, 2, 5}, Φ_3_ = {4, 6} and Φ_4_ = {7, 8, 9}.

#### 2.2.1 Partitioning the relative enzyme concentrations

When they are different co-regulation groups, and possibly singletons, the relative concentration of an enzyme can refer either to the sum of the enzyme concentrations of its group Φ_*q*_:

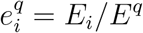

where 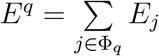, or to the total enzyme concentration in the pathway:

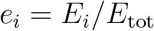

Hereafter, 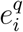 will be called the *intra-group relative concentration* and *e*_*i*_ will be called the *total relative concentration*.

The sum of enzyme concentrations in Φ_*q*_ relative to total enzyme concentration is:

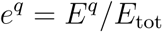

which will be called the *inter-group relative concentration*.

Therefore there is the following relationship between the three relative concentrations:

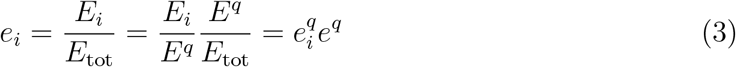

#### 2.2.2 Global co-regulation coefficient

Because *β*_*ij*_ ≠ 0 for enzymes of a given co-regulation group and *β*_*ij*_ = 0 for enzymes belonging to different co-regulation groups, the co-regulation matrix **M**_*β*_ can be separated into *p* submatrices, as shown in the example of table within figure 1. This makes it possible to specify the expression of the global co-regulation coefficient:

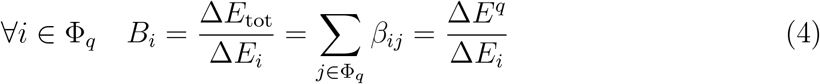

Thus, when there are different co-regulation groups, *B*_*i*_ quantifies also the impact due to co-regulations of a variation of *E*_*i*_ on the sum of enzyme concentrations of its group, which is the same as the impact on the total concentration.

If and only if *i* and *j* ∈ Φ_*q*_, we have the relationship:

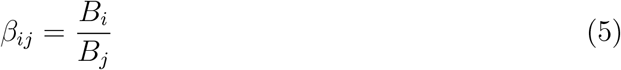

Note that 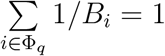 and 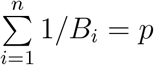.

#### 2.2.3 Parametric equation of the line of co-regulated enzyme concentrations

For a group Φ_*q*_ of co-regulated enzymes, the joint variation of concentrations reduces to one the number of degrees of freedom in this group. Therefore the intra-group relative concentrations move on a straight line in the space of intra-group relative concentrations. The parametric equation of this line, noted ℰ^*q*^, is (Coton et al. 2021):

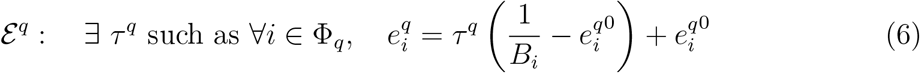

where *τ*^*q*^ is the driving variable and 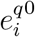 is the initial intra-group relative concentration of enzyme *i* in Φ_*q*_.

### 2.3 Evolution model

We studied the evolution of enzyme concentrations under selection for increased flux, assuming that the flux is proportional to fitness. We used the model developed by Coton et al. (2021), which is a differential equation system describing the variation of enzyme concentrations on a continuous timescale:

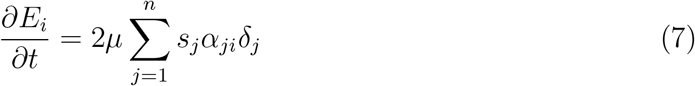

where:

- *µ* is the mutation rate
- 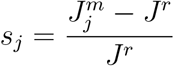 is the selection coefficient of a mutation targeting enzyme 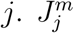 and *J*^*r*^ stand for the flux of the mutant and the flux of the resident, respectively.
- *δ*_*j*_ is the actual effect of the mutation targeting enzyme *j*
- *α*_*ji*_ is the redistribution coefficient measuring the impact on *E*_*i*_ of a mutation targeting enzyme *j*.

We used this differential system to study the evolution of enzyme concentrations when there are different groups of co-regulation in the pathway. We considered two cases, with and without competition for resources between all enzymes (respectively fixed *E*_tot_ and free *E*_tot_).

In both cases we searched for the evolutionary equilibria of relative enzyme concentrations, defined as (Coton et al. 2021):

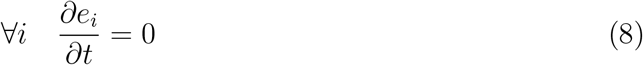

and we analysed the evolution of the flux over the generations of selection.

### 2.4 Computer simulations

The simulation method is the same as in Coton et al. (2021), albeit with different parameters. Briefly, we performed Markov Chain Monte-Carlo simulations using the R software (R Core Team 2017). We considered haploid populations of size *N* evolving in a constant environment (*X* = 1). At a given time step, all individuals in the population are identical, *i*.*e*. have the same phenotype *J* and the same enzyme concentrations (the “genotype”) **E** = (*E*_1_, *E*_2_, …, *E*_*n*_). Simulations were performed for pathways of different lengths (*n* = 3 to 10), with constant pseudo-activities **A** = (*A*_1_, *A*_2_, …, *A*_*n*_) and with co-regulation matrices **M**_*β*_ depending on the co-regulation structure of the pathway.

The vector of initial concentrations 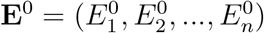 changed at each simulation. Initial concentrations 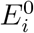 were drawn in a uniform law 𝒰 (0, 100) and redistributed proportionally to have 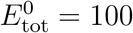. *E*_tot_ varied over time when there were no competition and was fixed at *E*_tot_ = 100 in the case of competition.

At each time step, a mutation targets randomly one of the enzymes and affects other enzymes according to the type of dependence considered. The values of the flux, of the selection coefficient and of the fixation probability of the mutant are computed. The mutation is fixed or not according to its fixation probability. Usually, the populations evolved over 125,000 time steps, and the system values were recorded every 250 steps.

These simulations allowed us to relax some of the assumptions of the mathematical analysis, namely small mutation effects, large population size and the absence of fixation of deleterious mutations.

Simulations were performed by using function `simul.evol.enz.multiple`.

### Data and code availability

Scripts of the simulations and custom functions, written in the R language (R Core Team 2017), are available in version 2 of the R package called `SimEvolEnzCons` (https://CRAN.R-project.org/package=SimEvolEnzCons).

## 3 Results

In a previous paper, we developed a model of joint evolution of enzyme concentrations under selection for increased flux under two cellular constraints, co-regulations between all enzyme concentrations and/or competition for resources (Coton et al. 2021).

Here, we generalize the model to the realistic case where different groups of coregulation coexist in a pathway. Within a group, the co-regulations can be positive (*positive groups*), or both positive and negative (*negative groups*). A group that contains only one enzyme is called a *singleton*. An enzyme in a singleton is independent from all other enzymes.

We searched for the evolutionary equilibrium of relative enzyme concentrations of the pathway, such as:

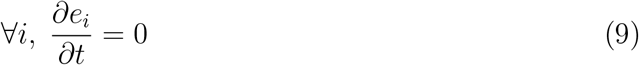

where *e*_*i*_ = *E*_*i*_*/E*_tot_ is the total relative concentration, with *E*_*i*_ the concentration of enzyme *i* and 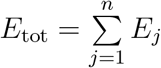 the total enzyme concentration.

We decomposed *e*_*i*_ into two parts: (i) the intra-group relative concentration 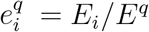, where *E*^*q*^ is the sum of enzyme concentrations in the co-regulation group Φ_*q*_; (ii) the inter-group relative concentration *e*^*q*^ = *E*^*q*^*/E*_tot_. These relative concentrations are linked by the relationship (see Material and Methods):

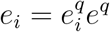

Taking account of these relative concentrations, equation 9 becomes:

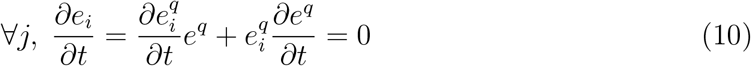

Therefore, solving 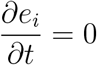 is equivalent to solve:

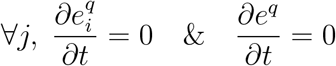

We ignore the other solution when 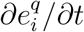 and *∂e*^*q*^/*∂t* are of opposite sign, because the equilibrium for total relative concentrations would probably be unstable.

Thus, the equilibrium of total relative enzyme concentrations implies both inter- and intra-group equilibria, that can be found separately.

Hereafter, we show how to find these equilibria and we analyse the flux variation over time. We considered two cases: without and with competition between all enzymes. Simulations of long-term evolution allowed us to test the validity of the analytical results.

### 3.1 Evolutionary equilibria without competition

The evolutionary equilibria within groups depend on the type of co-regulation between enzymes. We will consider positive groups, negative groups or both kinds of groups in the same pathway.

#### 3.1.1 Positive groups and singletons

##### 3.1.1.1 Intra-group equilibrium

The intra-group evolutionary equilibrium is defined by

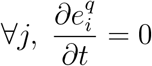

When the enzymes are all positively co-regulated, the solution of this equation is (Supporting Information II.3.1.2):

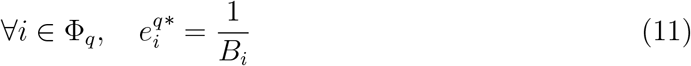

where *B*_*i*_ is the *global co-regulation coefficient*, which quantifies the impact due to coregulation of a variation of *E*_*i*_ on the sum of enzyme concentrations of group Φ_*q*_.

This *theoretical equilibrium* 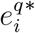 is achievable since *B*_*i*_ is positive for all *i*. If the group contains only one enzyme (*singleton*), we have 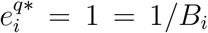 because *E*^*q*^ = *E*_*i*_ and *β*_*ii*_ = 1.

##### 3.1.1.2 Inter-group equilibrium

The inter-group evolutionary equilibrium is defined by

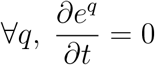

In Supporting Information II.3.2.2, we show that when all groups are positive or singletons, the solution of this equation is:

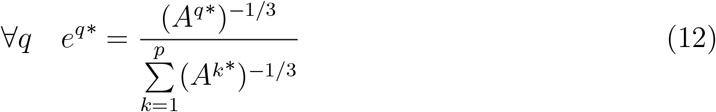

where *A*^*q*\*^, the *apparent activity of co-regulation group* Φ_*q*_, is a composite parameter defined such as:

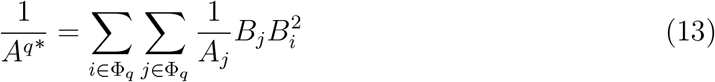

We see that the apparent activity of a group depends on the pseudo-activities of every enzyme in the group, weighed by the global co-regulation coefficients of this group. At the theoretical equilibrium, the inter-group relative enzyme concentrations *e*^*q*\*^ are inversely related to the *A*^*q*\*^.

##### 3.1.1.3 Total equilibrium

From equations 11 and 12, and using the relationship between relative concentrations 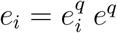 (equation 3), we obtain the theoretical equilibrium of the total relative concentrations when there are only positive groups or singletons:

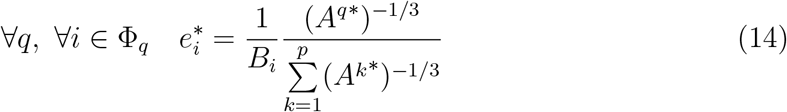

Note that if all enzymes are independent, we have *B*_*i*_ = 1 for all *i*, and we find the solution 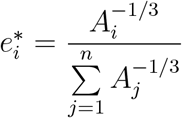 for all *i*; and if *all* enzymes are co-regulated, we have *p* = 1, and we find the solution 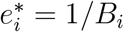 for all *i*. This is consistent with the results of Coton et al. (2021).

In the case of independence or positive co-regulation between enzymes, there is no limit to the increase of the flux over time.

#### 3.1.2 Negative groups

##### 3.1.2.1 Intra-group equilibrium

If there is at least one negative co-regulation in the group, at least one *B*_*i*_ is negative, which prevents the intra-group theoretical equilibrium above-mentioned (equation 11) to be reached since concentrations cannot be negative. There is another solution to find the intra-group equilibrium, based on the fact that when there is negative co-regulation, the flux cannot exceed a local maximum due the trade-off between enzyme concentrations in each negative group (Coton et al. 2021). At this point, the partial derivatives of the flux with respect to the concentrations cancel, so the flux response coefficient are all null in each negative group:

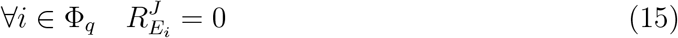

Using the expression of the flux response coefficient (equation 2), equation 15 becomes (Supporting Information II.3.1.3):

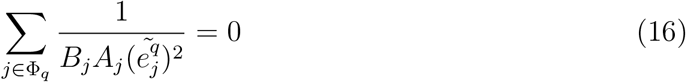

where 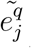 is called the *intra-group effective equilibrium*.

To solve this equation, we used the relationship between 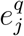 and *τ*^*q*^, the driving variable of the parametric equation of the line along which move the intra-group relative concentrations of co-regulated enzymes. From equation 6, equation 16 becomes:

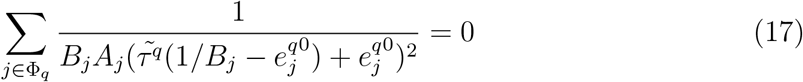

From this equation, 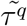, and therefore the intra-group effective equilibrium 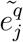, can be found numerically.

##### 3.1.2.2 Inter-group equilibrium

Equation 15 does not give a solution for the intergroup effective equilibrium. So to find *e*^*q*^ at equilibrium, we used the relationship between the sum *E*^*q*^ of enzyme concentrations in group Φ_*q*_ and the driving variable *τ*^*q*^ (Supporting Information II.3.3):

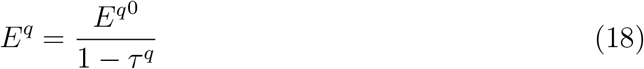

where 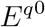 is the sum of initial enzyme concentrations in group Φ_*q*_.

If *all* groups are negative, the values 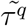 of the driving variables at the effective equilibrium (equation 17) allow us to compute the values of 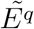 from equation 18. Thus we obtain the inter-group relative concentrations at effective equilibrium (Supporting Information II.3.2.3):

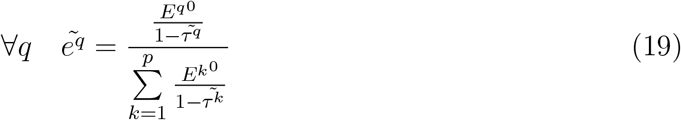

##### 3.1.2.3 Total equilibrium

Using equations 3, 16 and 19, the total effective equilibrium writes:

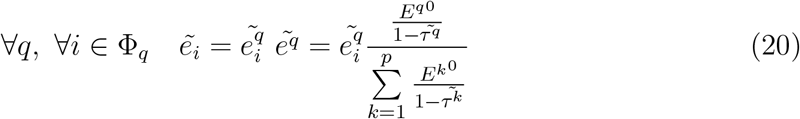

Due to the negative co-regulations between some enzyme concentrations, the flux cannot exceed a maximum. At this equilibrium, the flux reaches the following local maximum:

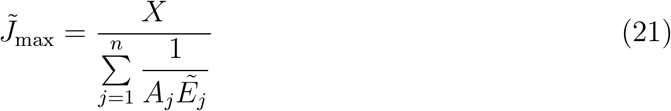

where (Supporting Information II.3.3):

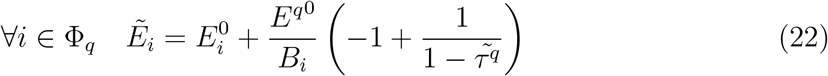

#### 3.1.3 Positive and negative groups

When positive (and/or singletons) and negative groups coexist in a pathway, the intragroup equilibria are reached within each group: the theoretical equilibrium 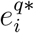 in positive groups and singletons (equation 11) and the effective equilibrium 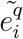 in negative groups (equation 16).

Regarding the *absolute* concentrations, *E*_*i*_ and *E*^*q*^ tends to infinity in the positive groups and singletons because no factor limits the increase of the *E*_*i*_’s, and respectively reaches 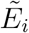 and 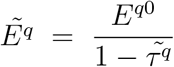 in the negative groups (equations 22 and 18). Therefore, over time, every 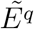 becomes negligible *relative to* the concentrations of positive groups/singletons and *E*_tot_ tends to 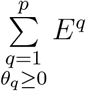. So, *e*^*q*^ (resp.*e*_*j*_) tends asymptotically to *e*^*q*^* (resp. 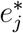) for positive groups and singletons, and asymptotically to 0 for negative groups.

Because *E*_*i*_ is limited to 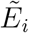 in the groups that reached their intra-group effective equilibrium, the flux tends asymptotically to a local maximum that writes (Supporting Information II.3.2.4):

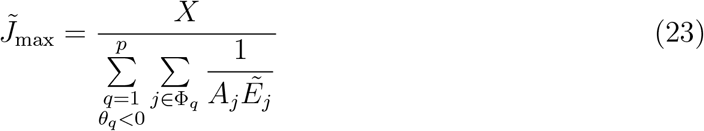

The different equilibria are summarized in Table 1.

**Table 1.**
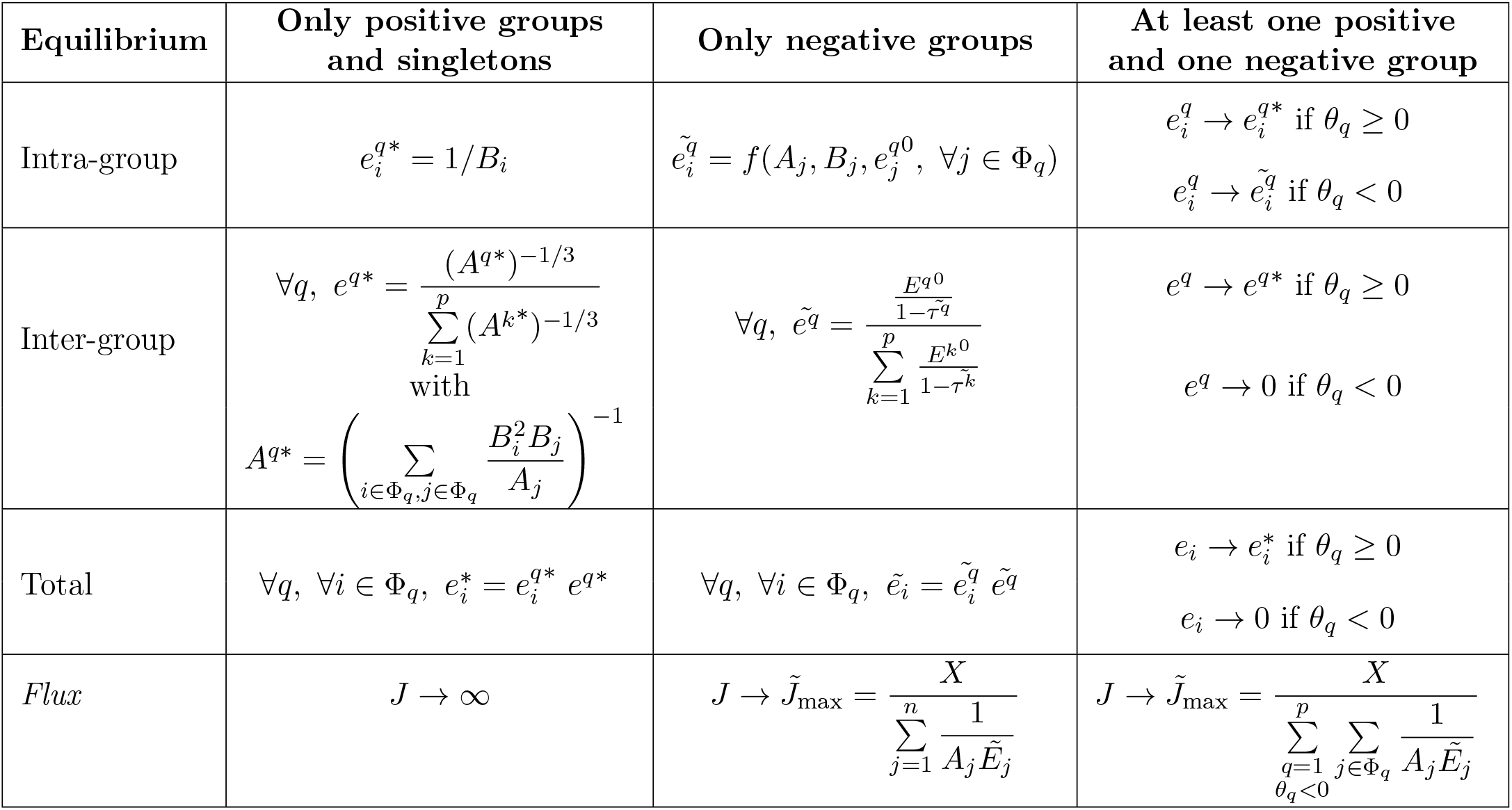
Equilibria of relative enzyme concentrations when there is no competition. *θ*_*q*_ is the indicator variable for the type of co-regulation in group Φ_*q*_ (see Material & Methods).

#### 3.1.4 Computer simulations

The simulations of long-term evolution for pathways of different lengths and with various types of co-regulation groups fully confirm the predictions:

- When there are only positive groups and/or singletons (figure 2A), the simulations show that the flux increases indefinitely, because there is no limit to the increase of enzyme concentrations. The total relative enzyme concentrations tend towards the predicted theoretical equilibrium, which depends on global co-regulation coefficients and enzyme pseudo-activities (figure 2B and 2C).

**Figure 2.**
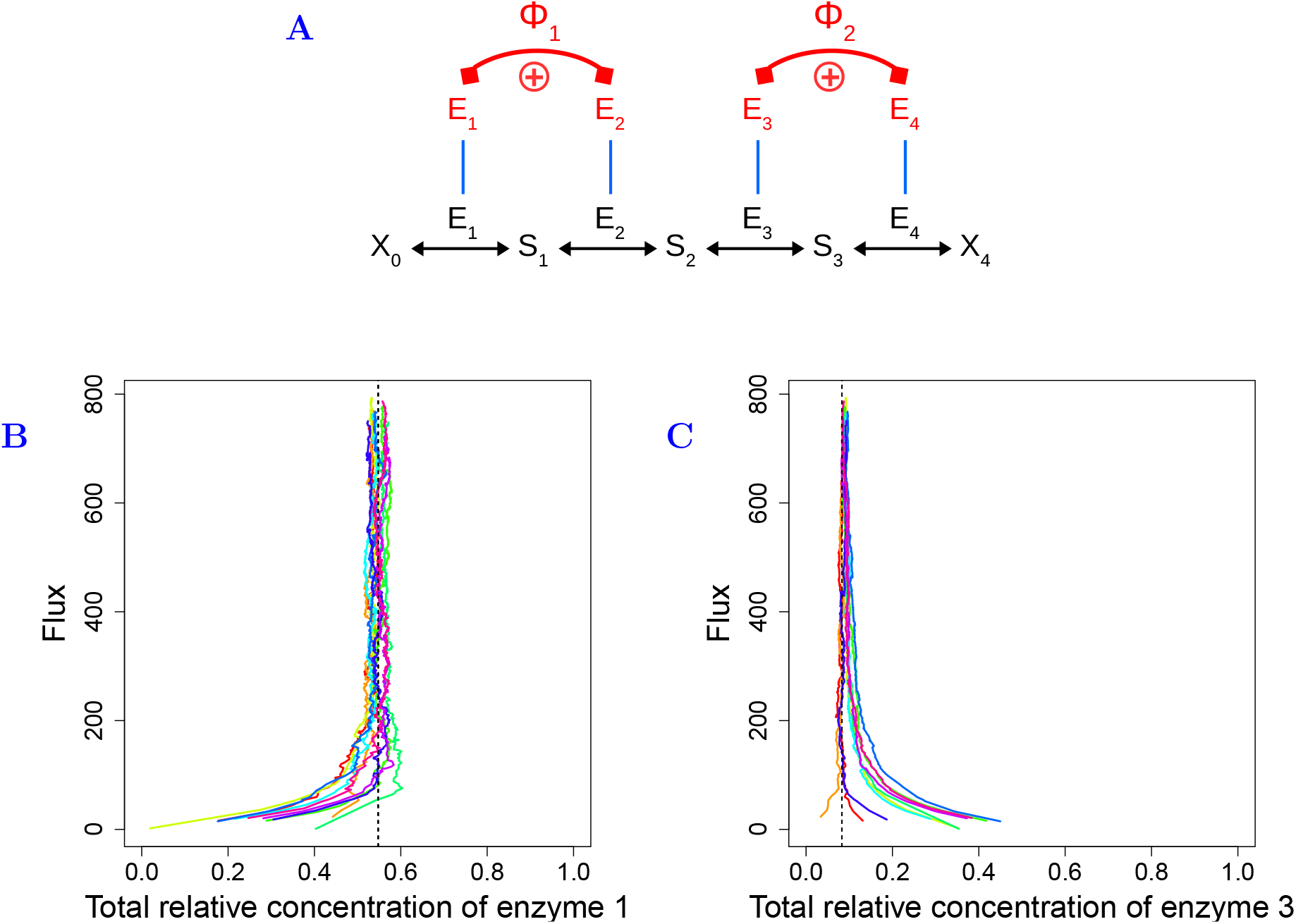
Simulations of enzyme evolution in a four-enzyme pathway when there are two positive groups, without competition. (A) The four-enzyme pathway used in the simulations. The enzymes E_1_ and E_2_ form a positive group and E_3_ and E_4_ form another positive group (same symbols as in figure 1). Parameter values: **A** = (1, 10, 30, 50), *X* = 1, *N* = 200, *β*_12_ = 0.32, *β*_43_ = 0.43. (B) Relationship between flux *J* and total relative concentration *e*_1_ of enzyme 1. The simulations run for 1 000 000 time steps. The results of 10 simulations are represented, each with specific initial concentrations (one color per simulation). Vertical dashed lines: abscissa of the theoretical equilibria of total relative concentrations 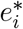. The total relative enzyme concentrations reach the theoretical equilibrium. (C) Same representation as in (B) for enzyme 3.
- When there are only negative groups (figure 3A), the *absolute* enzyme concentrations within each group are limited by the negative co-regulations. The total relative enzyme concentrations reach their effective equilibrium, which depends on enzyme pseudo-activities and on global co-regulation coefficients, but also on initial enzyme concentrations. Figures 3B and 3C show the joint evolution of total relative enzyme concentrations and flux. Whatever the initial concentrations, the flux increases until the maximum possible flux 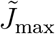 is reached, and enzyme concentrations get closer to their effective equilibrium. Then, enzyme concentrations fluctuate around the effective equilibrium while the flux stays approximately constant, which corresponds to the neutral zone.

**Figure 3.**
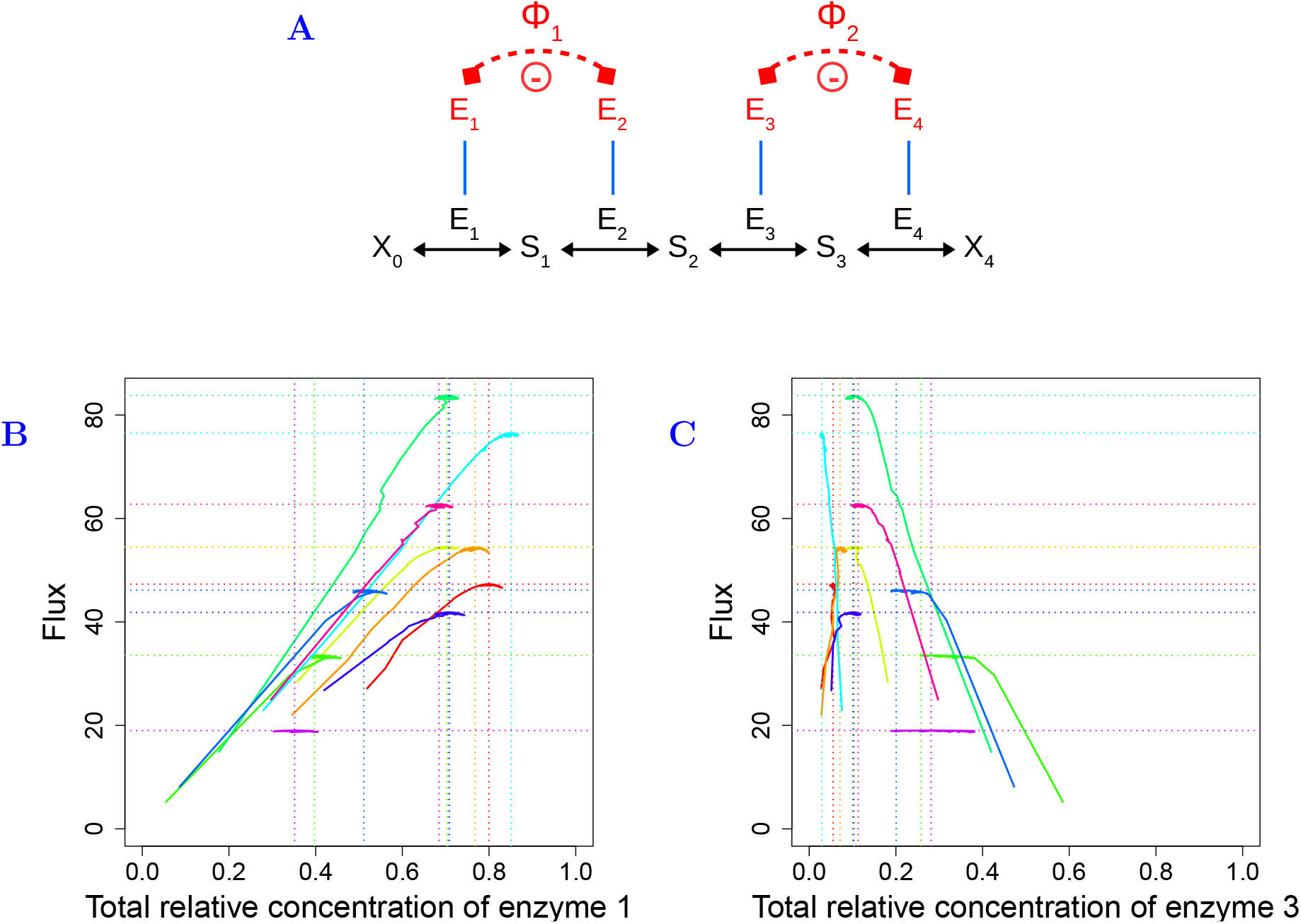
Simulations of enzyme evolution in a four-enzyme pathway when there are two negative groups, without competition. (A) The four-enzyme pathway used in the simulations. The enzymes E_1_ and E_2_ form a negative group and E_3_ and E_4_ form another negative group (same symbols as in figure 1). Parameter values: **A** = (1, 10, 30, 50), *X* = 1, *N* = 200, *β*_12_ = −0.1, *β*_43_ = −0.43. (B) Relationship between flux *J* and total relative concentration *e*_1_ of enzyme 1. The simulations run for 125 000 time steps. The results of 10 simulations are represented, each with specific initial concentrations (one color per simulation). Colored vertical dotted lines: abscissa of the effective equilibria of total relative concentrations 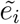. Colored horizontal dotted lines: maximal flux 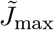. The total relative enzyme concentrations reach the effective equilibria, which correspond to the maximal flux. (C) Same representation as in (B) for enzyme 3.
- When positive (and/or singletons) and negative groups coexist (figure 4A), the intragroup relative concentrations are expected to reach the theoretical and effective equilibria, respectively. Due to the negative co-regulations, there is a maximum possible flux 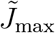 Figures 4B and 4C show the joint evolution of total relative enzyme concentrations and flux. We see that the flux gets closer and closer to 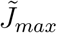 .After some burn-in period, total *relative* concentrations decrease in negative groups and increase in positive groups. However, predicted equilibria were never reached. Figures 4D and 4E show the joint evolution of absolute enzyme concentrations and flux in the same simulations as in figures 4B and 4C. After the same burn-in period, enzyme concentrations in positive groups increase slower and slower, while enzyme concentrations in negative groups stochastically move around the effective equilibria 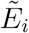 Indeed, as the flux gets closer to the maximum possible, selection intensity decreases and enzyme concentrations fluctuate within a quasi-neutral zone.

**Figure 4.**
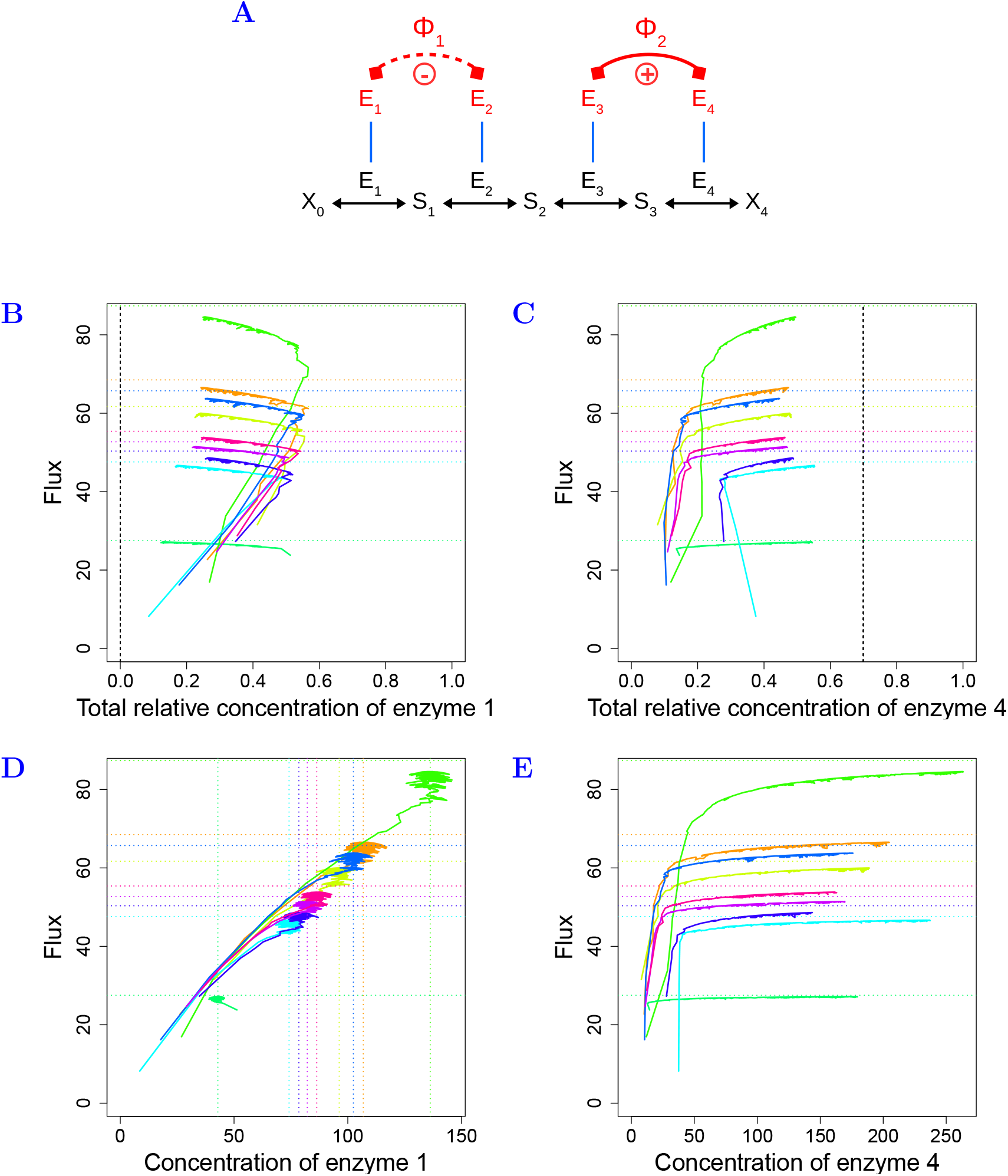
Simulations of enzyme evolution in a four-enzyme pathway when there are one positive and one negative groups, without competition. (A) The four-enzyme pathway used in the simulations. The enzymes E_1_ and E_2_ form a positive group and E_3_ and E_4_ form a negative group (same symbols as in figure 1). Parameter values: **A** = (1, 10, 30, 50), *X* = 1, *N* = 200, *β*_12_ = −0.32, *β*_43_ = 0.43. (B) Relationship between flux *J* and total relative concentration *e*_1_ of enzyme 1. The simulations run for 1 000 000 time steps. The results of 9 simulations are represented, each with specific initial concentrations (one color per simulation). Vertical dashed line: zero value, towards which the total relative enzyme concentrations of enzymes in the negative group tend. Horizontal dotted lines: maximal flux 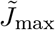. (C) Same representation as in (B) for enzyme 4. Vertical dashed lines: abscissa of the theoretical equilibria of total relative concentrations 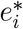. (D) Relationship between flux *J* and absolute concentration *E*_1_ of enzyme 1. Vertical dotted line: abscissa of the effective equilibria of absolute concentrations 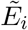. Horizontal dotted lines: maximal flux 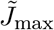. (E) Same representation as in (D) for enzyme 4.

Note that even though the stability of the equilibria could not be analytically proven, simulations show that the equilibria are stable.

### 3.2 Evolutionary equilibria with competition

Here we assume that there is competition for cellular resources between all enzymes of the pathway, so we fixed the total enzyme concentration. Thus the flux cannot exceed a maximal value whatever the type of co-regulation. The relationship between the flux and the enzyme concentrations is a dome in the multidimensional space of the concentrations. At the top of the dome the partial derivatives of the flux with respect to the concentrations cancel for all enzymes, so we have:

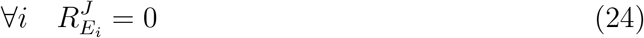

and at this point the enzyme concentrations are at their effective equilibrium.

In this section, we do not have to consider separately the positive and the negative groups, because competition between enzymes create negative relationships between all enzymes.

#### 3.2.1 Intra-group effective equilibrium

The intra-group relative concentration at effective equilibrium, 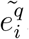, is solution of the following equation (Supporting Information II.4.1.2):

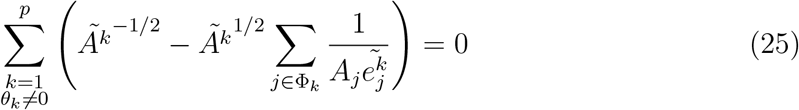

where the apparent activity *Ã*_*q*_ of group Φ_*q*_ at effective equilibrium is defined such as (provided *Ã*_*q*_ is positive):

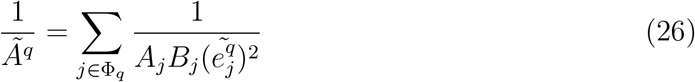

As previously, *Ã*_*q*_ is a weighed average of the pseudo-activities of enzymes, but its expression is different from the case without competition (equation 13).

We did not find any general analytical or numerical solutions for equation 25, but the simulations show that the equilibria exist (see section 3.2.4 below). We find a numerical solution in the particular case where there is only one co-regulation group – positive or negative – and any number of singletons. In that case, equation 25 writes after rearrangements (Supporting Information II.4.1.3):

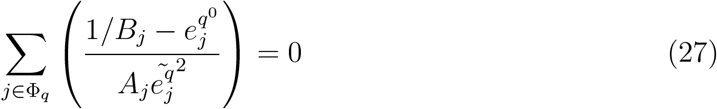

#### 3.2.2 Inter-group effective equilibrium

From equation 24, we can express 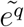, the inter-group relative concentration at effective equilibrium (Supporting Information II.4.1.1):

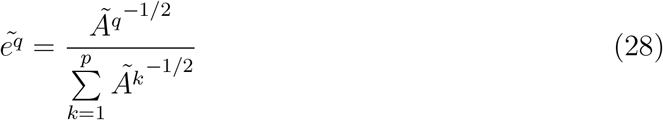

This expression is similar to that of the total relative concentrations at equilibrium when there is competition without co-regulations, except that the enzyme pseudo-activities are replaced by the apparent activities of co-regulation groups.

#### 3.2.3 Total effective equilibrium

From equations 3, 25 and 28 we get the total relative concentration 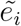 at effective equilibrium:

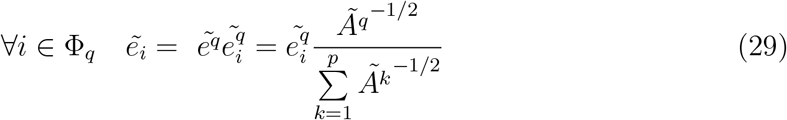

At this effective equilibrium, the flux is at a local maximum, 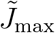, which is lower than the maximum flux *J*_max_ when there is no co-regulation because the co-regulations impose the flux to evolve in a lower dimension space.

The equilibria are summarized in table 2.

**Table 2.**
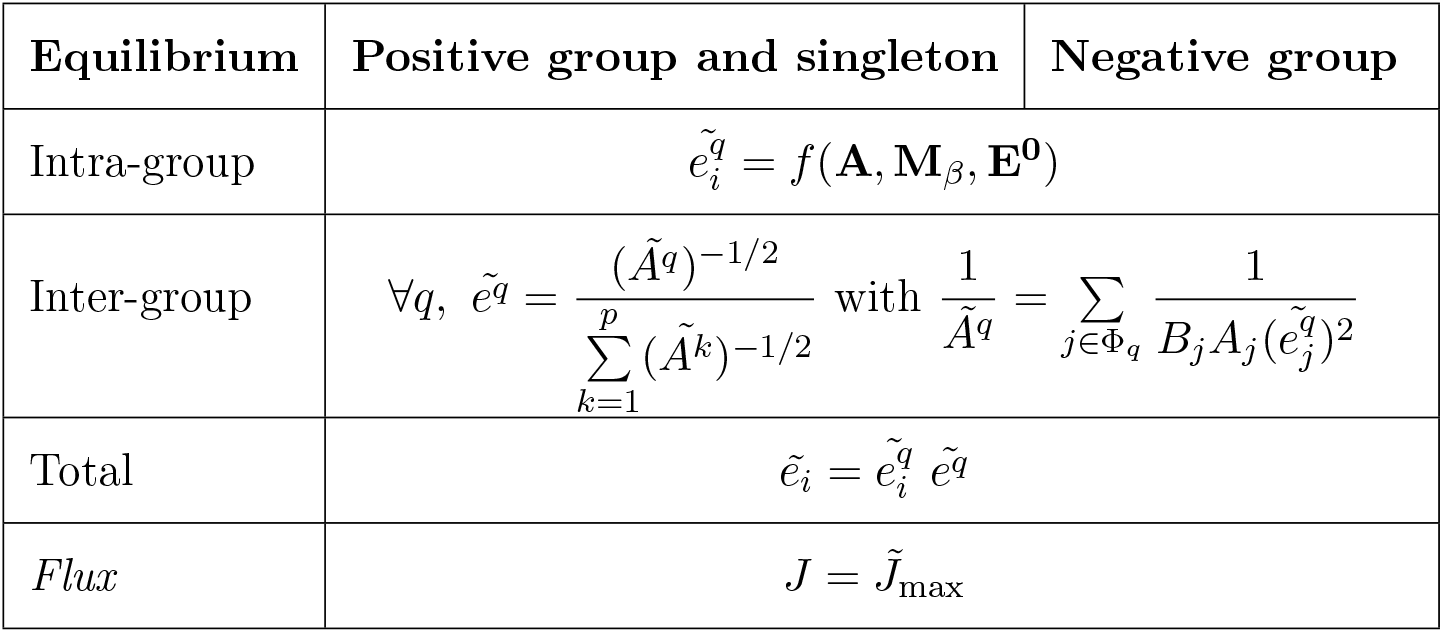
Effective equilibrium of relative enzyme concentrations when there is competition.

| Equilibrium | Positive group and singleton | Negative group |
| --- | --- | --- |
| Intra-group | $\tilde{e}_i^q = f(\mathbf{A}, \mathbf{M}_\beta, \mathbf{E}^0)$ | |
| Inter-group | $\forall q, \tilde{e}^q = \frac{(\tilde{A}^q)^{-1/2}}{\sum_{k=1}^p (\tilde{A}^k)^{-1/2}} \text{ with } \frac{1}{\tilde{A}^q} = \sum_{j \in \Phi_q} \frac{1}{B_j A_j (\tilde{e}_j^q)^2}$ | |
| Total | $\tilde{e}_i = \tilde{e}_i^q \tilde{e}^q$ | |
| Flux | $J = \tilde{J}_{\max}$ | |

#### 3.2.4 Computer simulations

Competition between enzymes creates negative relationships between all enzymes and the evolutionary issue is the same, whatever the co-regulation groups: given the initial conditions, the flux reaches a maximum, 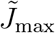, and total relative enzyme concentrations reach the effective equilibria. A numerical solution was found for the effective equilibria only in the case of a single co-regulation group and any number of singletons. Figure 5 shows an example of such situation. Figures 5B and 5C show the joint evolution of total relative enzyme concentrations and flux in the simulations. As predicted, the flux increases until a maximum, 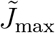, which depends on the initial conditions and that is bounded by the envelop-curve of the dome corresponding to the absence of co-regulation. Once 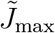 is reached, total relative enzyme concentrations fluctuate in a neutral zone around the effective equilibrium. Note that close to the equilibria, random fluctuations of enzyme-flux relationships delineate dome-shaped trajectories, which suggests that these trajectories evolve along a dome in lower dimension spaces as compared to the global competition dome. Another striking observation is that, before reaching the neutral zone, enzyme concentrations seem to walk erratically, sometimes decreasing before increasing again. Again, this suggests evolution in a constrained solution space.

**Figure 5.**
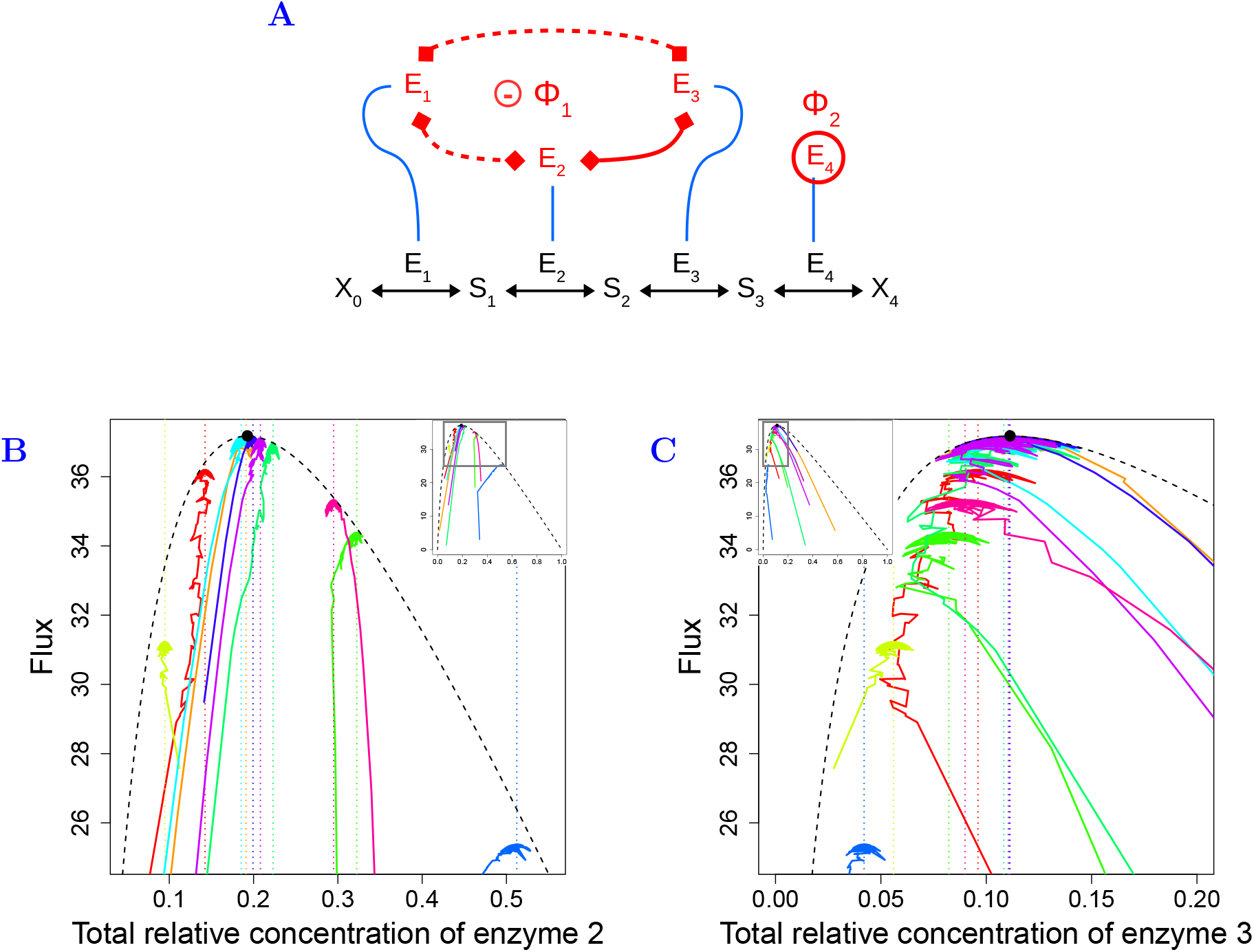
Simulation of enzyme evolution in a four-enzyme pathway with one co-regulation group and one singleton when there is competition. (A) Scheme of the four-enzyme pathway (same symbols as in figure 1). Enzymes E_1_, E_2_ and E_3_ form a negative group, and enzyme E_4_ is independent (singleton). The total concentration is fixed. Parameter values: **A** = (1, 10, 30, 50), *X* = 1, *N* = 200, *β*_12_ = 0.32, *β*_23_ = −0.43. (B) Relationship between flux *J* and total relative concentration *e*_2_ of enzyme 2. The simulations run for 125 000 time steps. The results of 10 simulations are represented, each with specific initial concentrations (one color per simulation). The main graph is a zoom of the frame drawn in the inset, which represents the entire graph. Colored vertical dotted lines: abscissa of the effective equilibria of total relative concentrations 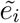. Black dashed curve: envelop curve of the dome when there is no co-regulations. All points are necessarily below this curve. Black point: summit of the envelop curve, which is the maximal flux *J*_max_ without co-regulations. (C) Same representation as in B for enzyme 3.

Figure 6A shows an example with one positive and one negative co-regulation group. In that case, numerical predictions for the effective equilibria are not computable. Figures 6B and 6C show the joint evolution of total relative enzyme concentrations and flux. Adaptive trajectories are qualitatively the same as in the previous case: given the initial conditions, a quasi-equilibrium is reached where the flux is close to a maximum, while total relative enzyme concentrations vary into a neutral zone around a fixed value.

**Figure 6.**
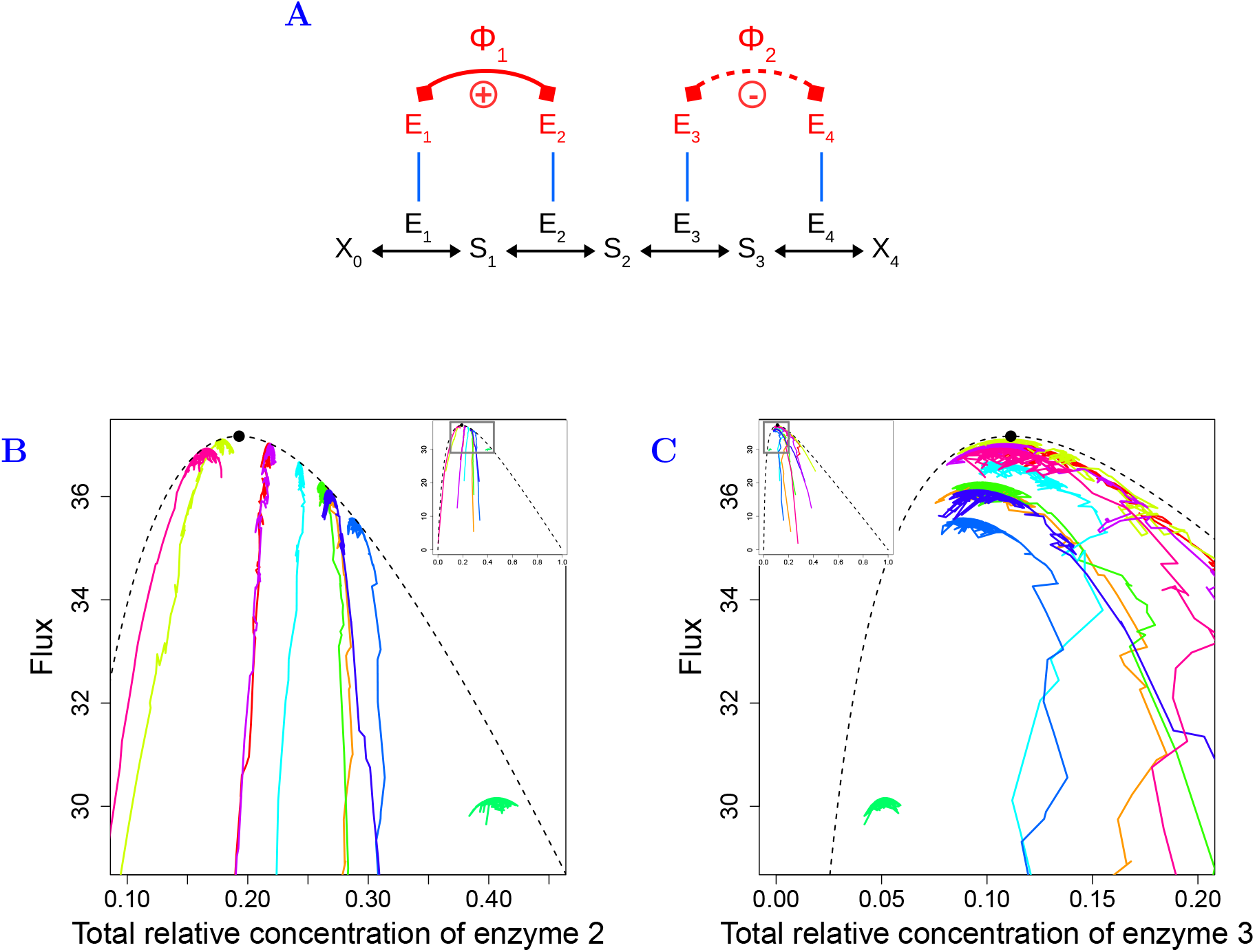
Simulation of enzyme evolution in a four-enzyme pathway with two co-regulation groups when there is competition. (A) Scheme of the four-enzyme pathway (same symbols as in figure 1). Enzymes E_1_ and E_2_ are positively co-regulated and enzymes E_3_ and E_4_ are negatively co-regulated. The total concentration is fixed. Parameter values: **A** = (1, 10, 30, 50), *X* = 1, *N* = 200, *β*_12_ = 0.32, *β*_43_ = −0.43. (B and C) Same representation as in (5B) and (5C) for enzymes 2 and 3 (the simulations run for 125 000 time steps). In this case, the effective equilibrium is not computable.

The dependency of the evolutionary equilibria to the initial conditions raises questions about the patterns of co-variation between enzyme concentrations that could be expected after independent bouts of evolution from different initial conditions. Figure 7 shows the relationships between the equilibrium concentrations of two enzymes belonging or not to the same co-regulation group for a four-enzyme pathway, starting from different initial conditions. In all cases where effective equilibrium were computable, we found a close adequacy between the predictions and the simulations (figures 7B, 7D). When all enzymes belong to the same co-regulation group (figure 7A), no relation was found (figure 7B). In all other cases (figures 7C, 7E, 7G), we found clear non-linear patterns (figures 7D, 7F, 7H). For example, in the case of a positive co-regulation group and a singleton, equilibrium concentrations of enzyme E_1_ and E_3_, which belong to the positive group, end on an ellipsoid (figure 7D). Other values of co-regulation coefficients give another ellipsoidal pattern (figures 7F, 7H). Without knowing the initial conditions, this shows that the constraint added by the co-regulations makes the evolution unpredictable with regards to enzyme concentrations.

**Figure 7.**
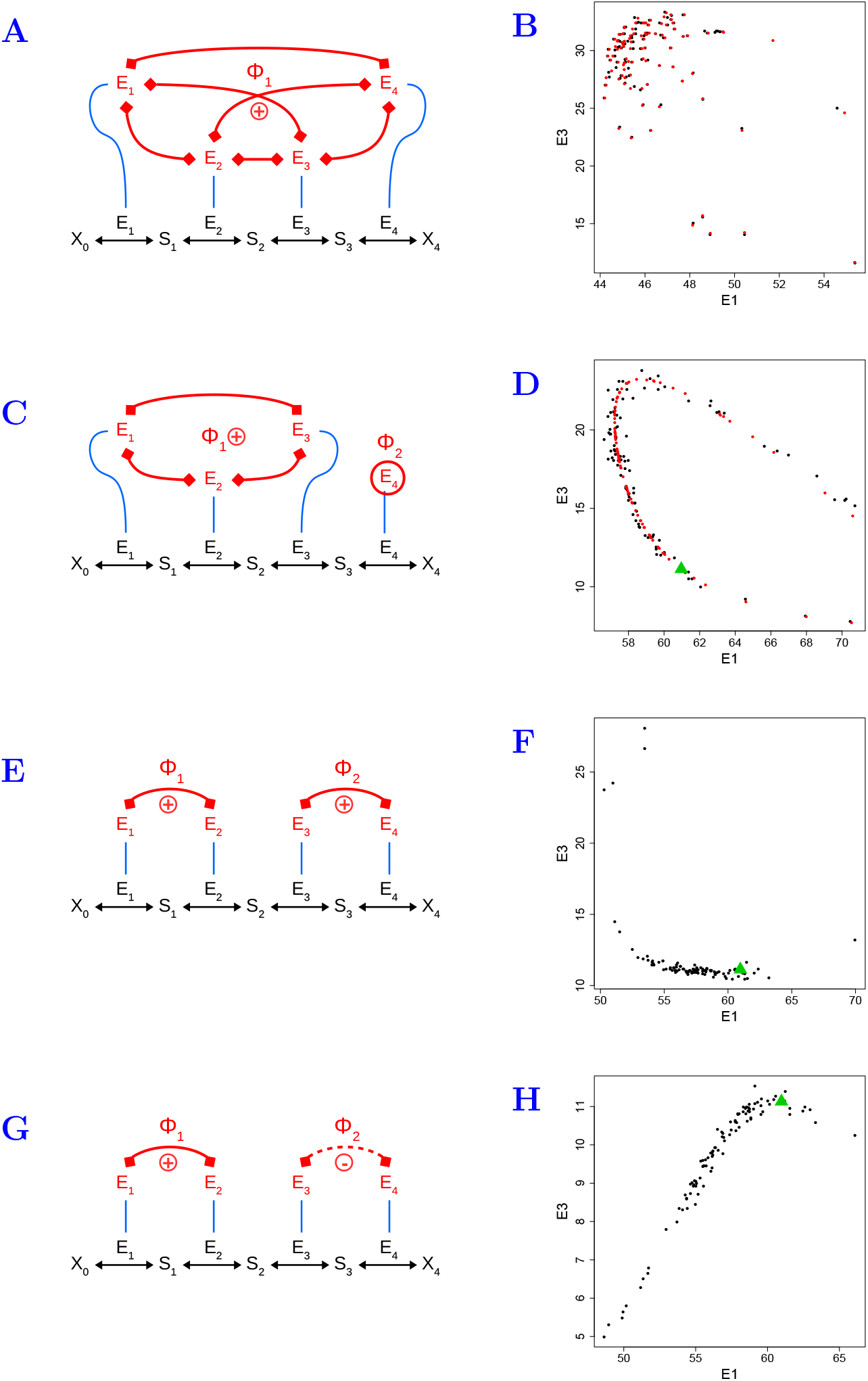
Simulations of evolution of enzyme concentrations for different types of co-regulation groups with competition. *Left column:* Different co-regulation groups in a four-enzyme pathway. Same symbols as in figure 1. Parameter values: **A** = (1, 10, 30, 50), *X* = 1, *N* = 10000. *Right column:* Relationship between the concentrations of enzymes E_1_ and E_3_ after 200 000 time steps. One hundred simulations were performed, which differed for the initial concentrations. Each black point corresponds to the end of one simulation. The red points correspond to the predicted effective equilibria, when computable. The green triangles indicate the enzyme concentrations at maximum flux when there is no co-regulation (*E*_1_ ≈ 61 and *E*_3_ ≈ 11.1). (A and B) All enzymes are positively co-regulated (*β*_12_ = 0.32, *β*_23_ = 2, *β*_34_ = 0.1). The simulated and predicted effective equilibria are close to each other, which confirms the results of our analysis. (C and D) Enzymes E_1_, E_2_ and E_3_ are positively co-regulated and enzyme E_4_ is independent (*β*_12_ = 0.1, *β*_23_ = 2). The simulated effective equilibria are close to the curve described by the predicted effective equilibria. (E and F) There are two groups of positively co-regulated enzymes, E_1_ and E_2_ on the one hand and E_3_ and E_4_ on the other hand (*β*_12_ = 0.32, *β*_34_ = 0.1). We hypothesize that the curve described by the black points follow the curve of effective equilibria that we could not compute. (G and H) The enzymes E_1_ and E_2_ are positively co-regulated and enzymes E_3_ and E_4_ are negatively co-regulated (*β*_12_ = 0.32, *β*_43_ = −0.43). The black points also describe a curve, but unlike the previous case, the relationship between *E*_1_ and *E*_3_ is positive.

### 3.3 The range of selective neutrality of enzyme concentrations

The ranges of neutral variation (RNV) of enzyme concentrations close to the equilibria were compared in the different situations.

#### 3.3.1 RNV when there is no competition

Three cases can be distinguished, as we did for the search of equilibria:

1. When there are only positive groups and singletons, the flux response coefficients 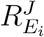 are constant at theoretical equilibrium (Supporting Information II.5). Therefore, the selection coefficient *s*_*i*_, through the relationship 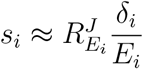 (Coton et al. 2021), tends to zero because *E*_*i*_ increases indefinitely. The mutations have less and less effect on the flux while approaching the theoretical equilibrium.
2. When there are only negative groups, the flux response coefficients, and hence the selection coefficients, are null at effective equilibrium, so there is selective neutrality.
3. When there are both positive and negative groups, the flux response coefficients are null for negative groups at effective equilibrium and tends to zero for positive and singleton groups (Supporting Information II.5). Therefore the selection coefficients tend to zero in every group.

Thus, when there is no competition, there is neutrality close to the equilibria, whatever the number and type of co-regulation groups. However, unlike what we had gone before Coton et al. (2021), we did not succeed in determining the limits of the RNV due to the complexity of the relationships between the co-regulation groups.

In the simulations, the neutral zone is revealed by the fluctuations of the concentrations around the effective equilibrium for negative groups (figures 3 and 4D). Figures 3B and 3C also shows that the range of neutral variation (RNV) of enzyme concentrations depend both on the initial concentrations and on the enzyme parameters.

#### 3.3.2 RNV when there is competition

When there is competition, the response coefficients are null at effective equilibrium, and so are the selection coefficients. Therefore, there is a neutral zone around every enzyme concentration at equilibrium. However, as previously, we could not untangle the factors that determine the RNV. In the simulations, the neutral zone is clearly visible (figures 5 and 6). At the end of the simulations, the concentrations fluctuate around the predicted effective equilibria, making apparent the ranges of neutral variation at the top of domes. At equilibrium, RNVs probably depend on pseudo-activities, co-regulation coefficients and initial concentrations.

## 4 Discussion

The enzyme-flux relationship is commonly used to model the genotype-phenotype relationship, owing to the sound theoretical framework developed for decades that link enzymes properties (the “genotype”) to the flux (the phenotype) (Wright 1934; Kacser and Burns 1981; Fiévet et al. 2018). In the framework of evolutionary theory, this multidimensional relationship constitutes an adaptive landscape when the flux is proportional to fitness.

In Coton et al. (2021), we developed a model of evolution of enzyme concentrations under selection for increased flux. We applied two constraints, competition for resources and/or co-regulation between all enzymes, and showed the existence of two types of evolutionary equilibrium of relative concentrations, depending on the presence of constraints or not. However, considering that all enzymes are co-regulated in a pathway is a particular case. Here, we analysed the general case where there are different co-regulation groups, with or without competition, distinguishing two types of co-regulation groups: (i) the positive groups when enzymes are all positively co-regulated; (ii) the negative groups when there is at least one negative co-regulation in the group. We called singletons the enzymes that are independent of all other enzymes.

To predict the evolutionary equilibria, we decomposed the total enzyme relative concentrations into intra- and inter-group relative concentrations. In most situations, analytical or numerical solutions were found and were confronted to the results of computer simulations. Combining theoretical predictions and simulation results, we showed that a stable evolutionary equilibrium for relative enzyme concentrations always exists. In the trivial case of only positive regulation groups without competition, this equilibrium is unique whatever the initial concentrations. In all other cases, evolutionary equilibria depended not only on enzyme parameters and on the gene regulatory network – through the co-regulation matrix –, but also on initial conditions. Such an evolutionary contingency emerges from the existence of constraints, which stem either from the competition for cellular resources that limits total enzyme concentration, or from negative co-regulations, that prevents the simultaneous increase of the concentrations of the co-regulated enzymes.

### 4.1 One negative co-regulation is sufficient to limit the flux

Even in the absence of competition, we showed that the trade-offs between co-regulated enzymes in negative groups blocks the flux at a local maximum and are sufficient to limit the evolution of all enzyme concentrations. Heckmann et al. (2018) observed a similar phenomenon. They simulated the evolution of enzyme turnovers (*k*_cat_) under selection for increased growth rate with fixed concentrations. Because enzyme concentrations and turnover numbers have concurrent roles in the expression of the flux (equation 1), constraining enzyme concentrations is similar to fixing enzyme turnovers, as we did here. Fixing some *k*_cat_’s, Heckmann et al. (2018) showed that the diminishing returns in the relationship between the variable *k*_cat_’s and growth rates prevented selection for high *k*_cat_’s and maintained growth rate far from its possible maximum value. Here, we showed that any biological constraint that prevents the concentration of one enzyme from increasing despite selection for increasing the flux will result into an upper-bound for the flux at the evolutionary equilibria. Other constraints could stem from trade-offs between active sites of an enzyme (Savir et al. 2010), or from the gene regulatory network. For example, among the genes that control yeast cell cycle, the cyclins Clb1 and Clb2 deactivate different transcription factors like MBF, SBF and Swi5 that in turn control different parts of the cycle (Li et al. 2004).

Another interesting result is that upon selection, the flux increases monotonically towards the maximum, whereas enzyme concentrations decrease or increase, depending on the target of the mutations and on their effect on the flux *via* the co-regulated enzymes (figures 5 and 6). Thus, a monotonous phenotypic response to directional selection does not necessarily mean that the underlying genotypic factors evolve monotonously.

### 4.2 Evolutionary contingency and the world of possibles

A major consequence of the existence of co-regulations is that both maximum flux and evolutionary equilibria for total relative enzyme concentrations depend on initial conditions. In this model, the outcome of the evolutionary process depends on history. The knowledge of the gene regulatory network and enzyme parameters is not sufficient to predict the evolutionary outcome. At most, it would be possible to predict an envelop-curve for the flux in the competition case (figures 5 and 6).

The dependence on initial conditions also results in non linear patterns between enzyme concentrations at evolutionary equilibria (figure 7). This result could explain the wide range of genetic variation of enzyme concentrations observed within populations, as well as their highly polygenic nature (Damerval et al. 1994; Albert and Kruglyak 2015; Chick et al. 2016). Such dependency to initial conditions is often observed in complex biological systems. It is the case for non linear dynamical systems like the lactose operon (Laurent and Kellershohn 1999), but also for complex patterns emerging from network self-organization that can be treated using game theory (Broere et al. 2017).

Contingency comes from the fact that constraints modify the adaptive landscape by limiting the ability to move in the multidimensional space (**E**, *J*). We searched the number of degrees of freedom of the system, which is the number of independent variables necessary and sufficient to describe the system, given the constraints applied and the fixed parameters – here the pseudo-activities **A**, the co-regulation coefficients of matrix **M**_*β*_ and the initial concentrations **E**^**0**^ (Supporting Information III.1).

If there is no constraint on enzyme concentrations, the genotypes move freely in the multidimensional space (**E**, *J*). When there is competition between all enzymes, the adaptive landscape becomes a dome in (**E**, *J*), the shape of which depends only on the pseudo-activities (Coton et al. 2021). When there are co-regulations between *all* enzymes, the enzyme concentrations are linearly related and are fully determined by the driving variable *τ* : the adaptive landscape is reduced to two dimensions, *τ* and *J* (Coton et al. 2021). It is the same when we add competition. In that case the adaptive landscape becomes a section of the dome, oriented by the global co-regulation coefficients and the initial concentrations.

Here we considered the presence of co-regulation groups. Without competition, the enzyme concentrations are linearly related within groups, and are independent from enzyme concentrations of other groups. So the concentrations of each co-regulation group *q* are given by its driving variable *τ*^*q*^. The adaptive landscape is in a *p* + 1 dimension space – the number of co-regulation groups plus the flux. With competition between all enzymes, we could not determine precisely the number of degrees of freedom. Simulations showed that the adaptive landscape is a projection in lower dimensions of the dome due to competition, and its shape is also a dome (figure 6). The effective equilibria are given by the projection of the top of the new dome in the concentration space. They depend on the number of co-regulation groups, and each additional co-regulation between two previously independent groups decreases the number of dimensions.

### 4.3 Evolutionary jumps and selective neutrality

Generalizing the results of Hartl et al. (1985) on the “natural selection of selective neutrality”, we showed that in all cases there is a neutral zone of enzyme concentrations, where small changes in concentrations do not significantly change the flux that is the target of selection. The ranges of neutral variations differ according to the pseudo-activities, the co-regulation coefficients and the initial conditions.

In our simulations, we found that, given the initial conditions, the system evolved quite rapidly to the equilibrium. In the same way, any mutational change in a pseudo-activity or a co-regulation coefficient would modify the adaptive landscape and change simultaneously enzyme concentrations and flux, defining new equilibria. Such evolutionary jumps, also mentioned by Heckmann et al. (2018) when mutations affect multifunctional enzymes, are reminiscent of punctuated equilibria (Gould and Eldredge 1977). Altogether, our simplified model of enzyme evolution allowed us to give support to observations in evolution: the selective neutrality of enzymes at evolutionary equilibria, and the existence of evolutionary jumps due to rare mutations changing system’s parameters.

## Acknowledgements

CC was supported by a PhD thesis grant from the French Ministère de l’Enseignement Supérieur, de la Recherche et de l’Innovation.

The authors declare no conflict of interest.

## Author contributions

CD and DdV initiated the project. CC developed the model, performed the analysis and the simulations and developed the dedicated R package. CC and DdV wrote the manuscript, with the contribution of CD.

## Appendix A

### Glossary of mathematical symbol

#### Latin symbols

*A*_*i*_: Pseudo-activity of enzyme *i*, a composite parameter including the kinetic properties (catalytic constant and Michaelis-Menten constant) of the enzyme and thermodynamic equilibrium of upstream reactions
A: Vector of enzyme pseudo-activities
*A*^*q*\*^: Apparent activity of group Φ_*q*_ at theoretical equilibrium without competition
*Ã*_*q*_: Apparent activity of group Φ_*q*_ at effective equilibrium with competition
*B*_*i*_: Global co-regulation coefficient, representing effect of a variation of *E*_*i*_ on *E*_tot_ due to co-regulation, and also of *E*_*i*_ on *E*^*q*^
*E*_*i*_: Concentration of enzyme *i*
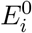: Initial concentration of enzyme *i*
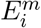: Concentration of a mutant enzyme *i*
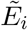: Concentration of enzyme *i* at effective equilibrium
E: Vector of enzyme concentrations, considered as the genotype
*E*^*q*^: Sum of enzyme concentrations in Φ_*q*_
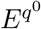: Initial sum of enzyme concentrations in Φ_*q*_
*E*_tot_: Total enzyme concentration in the metabolic pathway
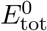: Initial total enzyme concentration
*e*_*i*_: Total relative concentration of enzyme *i*; *e*_*i*_ = *E*_*i*_*/E*_tot_
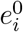: Initial total relative concentration of enzyme *i*
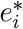: Total relative concentration of enzyme *i* at theoretical equilibrium
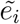: Total relative concentration of enzyme *i* at effective equilibrium
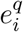: Intra-group relative concentration of enzyme *i* in Φ_*q*_ ; 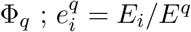
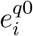: Initial intra-group relative concentration of enzyme *i* in Φ_*q*_
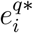: Intra-group relative concentration of enzyme *i* in Φ_*q*_ at theoretical equilibrium
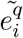: Intra-group relative concentration of enzyme *i* in Φ_*q*_ at effective equilibrium
*e*^*q*^: Inter-group relative concentration of enzyme of Φ_*q*_ in regard to total concentration; *e*^*q*^ = *E*^*q*^*/E*_tot_
*e*^*q*\*^: Inter-group relative concentration of enzyme *i* in Φ_*q*_ at theoretical equilibrium
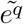: Inter-group relative concentration of enzyme *i* in Φ_*q*_ at effective equilibrium
ℰ ^*q*^: Line in the space of intra-group relative enzyme concentrations along which the intra- group relative concentrations vary in the case of co-regulation for each group *q*
*J*: Metabolic flux, considered as a phenotype that is proportional to fitness
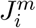: Flux resulting from a mutation targeting enzyme *i*
*J*^*r*^: Flux of the resident
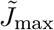: Maximal value of the flux, corresponding to the effective equilibrium
**M**_*β*_: Matrix of co-regulation coefficients
*m*_*q*_: Number of enzymes in Φ_*q*_
*N*: Effective population size
*n*: Number of enzymes in the metabolic pathway
*p*: Total number of co-regulation groups
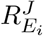: Flux response coefficient of enzyme *i*
S_*i*_: Metabolite *i*, product of reaction *i* and substrate of reaction *i* + 1
*s*_*i*_: Selection coefficient of a mutation targeting enzyme *i*
*X*: Constant representing the environment, depending on the concentrations of the input substrate X_0_ and the output product X_*n*_ of the pathway

#### Greek symbols

*α*_*ij*_: Redistribution coefficient, expressing the variation of *E*_*j*_ due to a mutation targeting *E*_*i*_
*β*_*ij*_: Co-regulation coefficient of enzyme *i* on *j*, expressing the change of *E*_*j*_ due to co-regulation when *E*_*i*_ varies
*δ*_*i*_: Actual effect of a mutation affecting enzyme *i*
*θ*_*q*_: Indicator variable indicating the type of co-regulation in the co-regulation group *q*
*µ*: Mutation rate at each time unit
*ν*: Canonical effect of a mutation
*τ*^*q*^: Driving variable of the line ℰ ^*q*^ in the space of intra-group relative concentrations for enzymes in the group Φ_*q*_
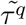: Value of the driving variable at the effective equilibrium for the group Φ_*q*_
Φ_*q*_: Co-regulation group *q*

#### Other symbols

0: Exponent that refers to the initial state of the system
*: Exponent that refers to the theoretical equilibrium
∼: Refers to the effective equilibrium

