## Supporting Information for "Evolution of enzyme levels in metabolic pathways: A theoretical approach. Part 2"

Charlotte Coton, Christine Dillmann, Dominique de Vienne

November 18, 2021

### I Supplementary Figures

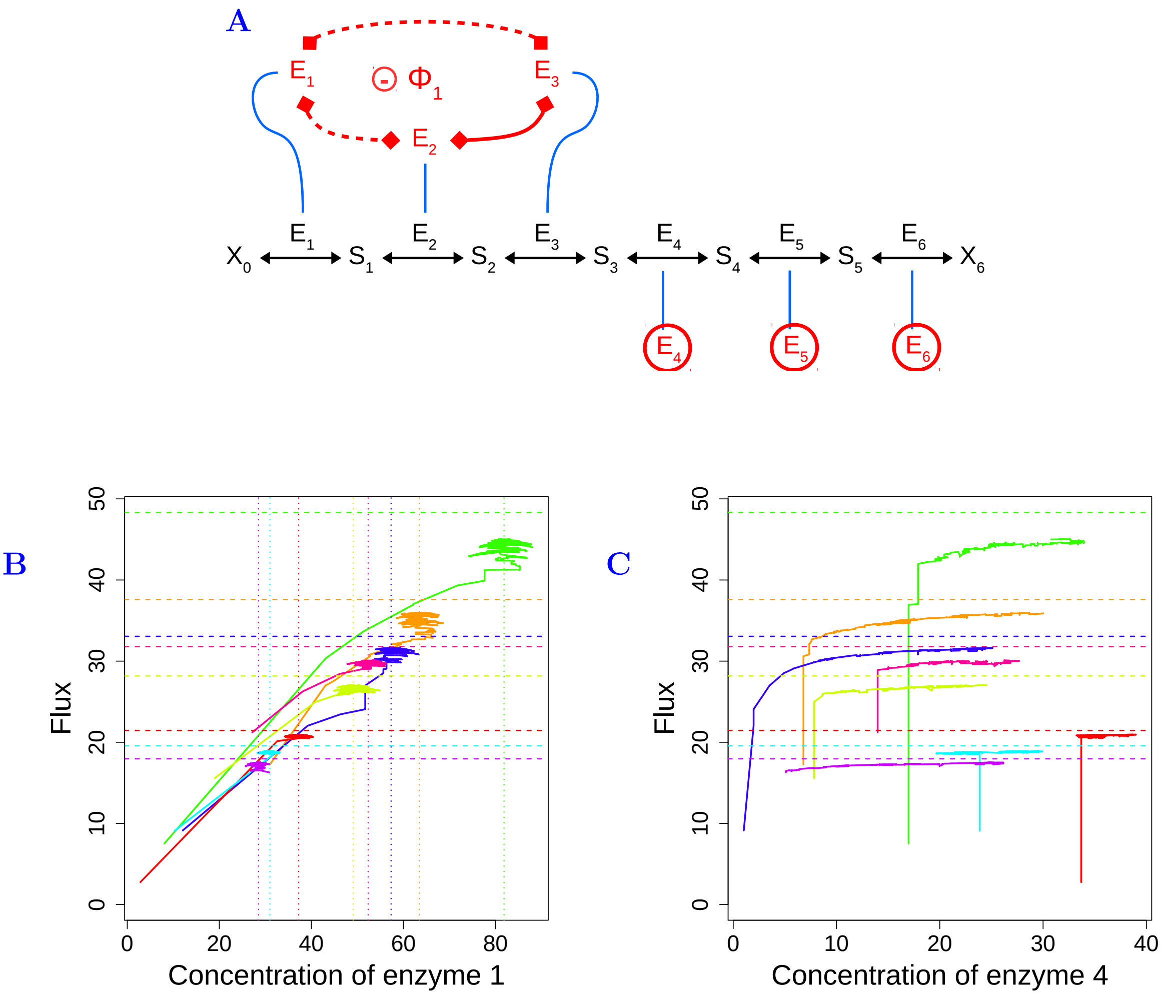

**Figure S1. Simulations of enzyme evolution in a six-enzyme pathway when there is a negative co-regulation group and three singletons, without competition.** (A) The six-enzyme pathway used in the simulations. The first three enzymes $E_1$, $E_2$ and $E_3$ form a negative group and the other enzymes are independent (same symbols as in figure 1). Parameter values: $\mathbf{A} = (1, 10, 30, 50, 100, 200)$, $X = 1$, $N = 200$, $\beta_{12} = -0.32$, $\beta_{23} = 0.43$. (B) Relationship between flux $J$ and concentration of enzyme 1. The simulations run for 125 000 time steps. The results of 8 simulations are represented, each one with specific initial concentrations (one color per simulation). Vertical dotted lines: abscissa of the effective equilibria of absolute concentrations $\tilde{E}_i$. Horizontal dashed lines: maximal flux $\tilde{J}_{\max}$. The concentrations of the negatively co-regulated enzymes first increase, then stabilize around their effective equilibrium, which prevents the flux from increasing beyond a local maximum. (C) Same representation as in (B) for independent enzyme 4. Even if there is no theoretical limit, the concentration of the independent enzyme tends towards a plateau because there is no more selective pressure when $\tilde{J}_{\max}$ is reached.

**A**

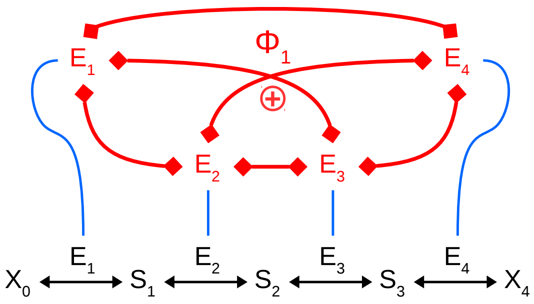

**B**

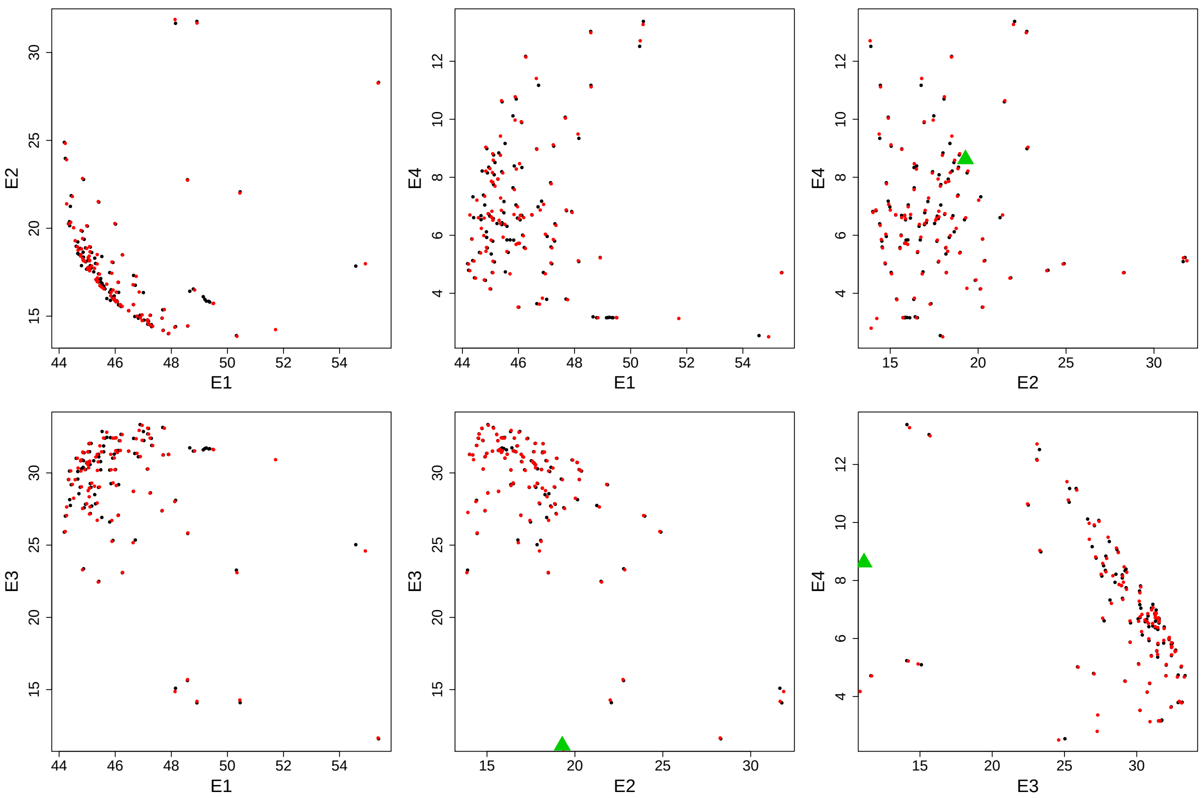

**Figure S2. End of simulation of enzyme evolution in a four-enzyme pathway when all enzymes are co-regulated with competition.** (A) The four-enzyme pathway used in these simulations. All enzymes are positively co-regulated. Parameter values: $\mathbf{A} = (1, 10, 30, 50)$, $X = 1$, $N = 10000$, $\beta_{12} = 0.32$, $\beta_{23} = 2$ $\beta_{34} = 0.1$. (B) Relationship between the enzymes concentrations at the end of simulation. One hundred simulations were performed, with different initial concentrations. Each black point correspond to the end of one simulation. The green triangle ($E_1 = 61$, $E_2 = 19.3$, $E_3 = 11.1$, $E_4 = 8.6$) corresponds to the maximum flux without regulation. The black points seem to be random, but actually depends on the position of 1/B and $E^0$ is the 4D-dome.

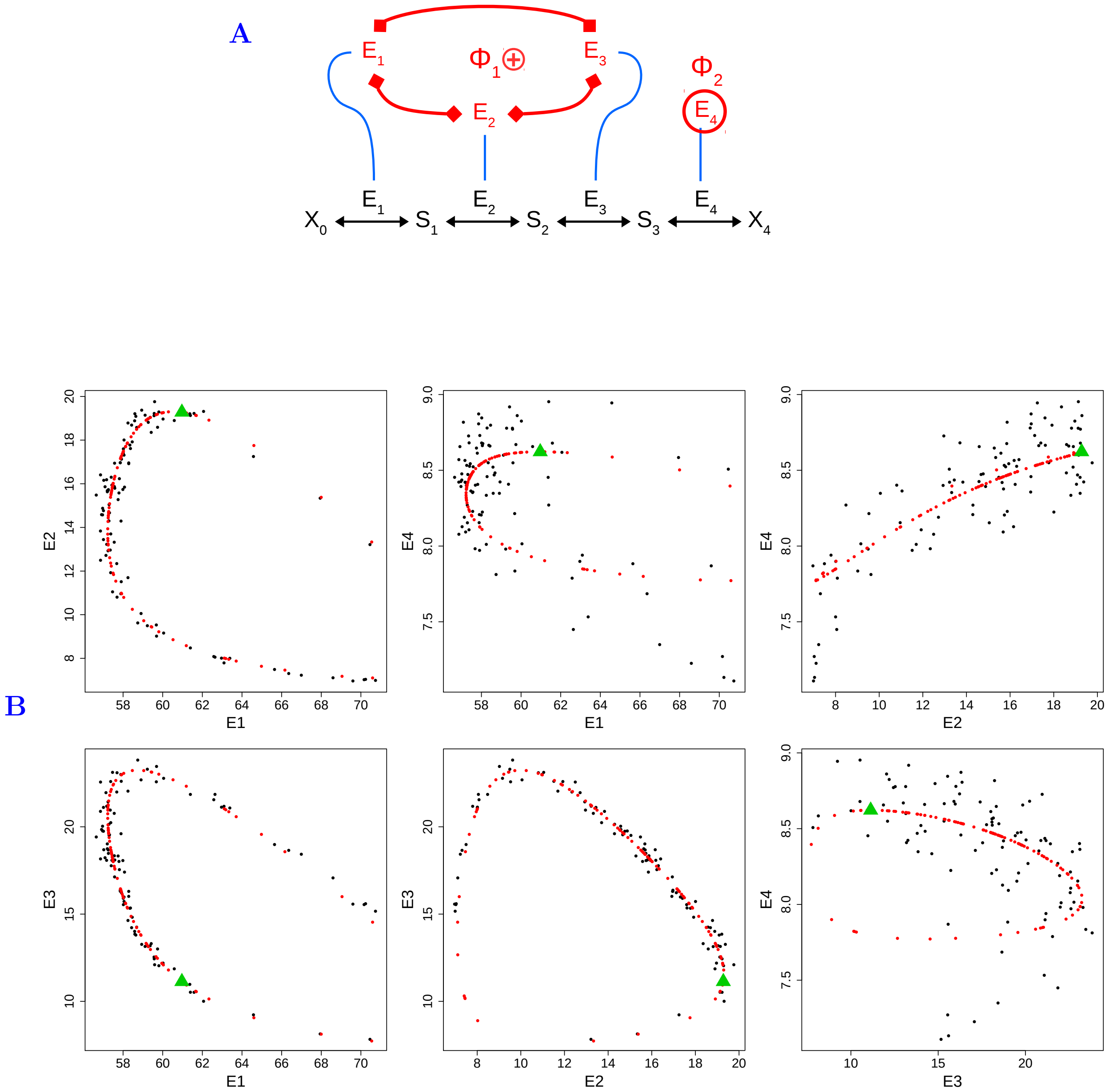

**Figure S3. End of simulation of enzyme evolution in a four-enzyme pathway when there is one positive group and one independent enzyme with competition.** (A) The four-enzyme pathway used in these simulations. The enzymes $E_1$, $E_2$ and $E_3$ are positively co-regulated, whereas enzyme $E_4$ is independent. Parameter values: $\mathbf{A} = (1, 10, 30, 50)$, $X = 1$, $N = 10000$, $\beta_{12} = 0.1$, $\beta_{23} = 2$. (B) Relationship between the enzymes concentrations at the end of simulation. One hundred simulations were performed, with different initial concentrations. Each black point correspond to the end of one simulation. The red points correspond to the predicted effective equilibrium. The green triangles ($E_1 = 61$, $E_2 = 19.3$, $E_3 = 11.1$, $E_4 = 8.6$) correspond to the maximum flux without regulation. The black points describe a curve close to the curve of predicted equilibria in each dimension. The average distance between simulated and predicted equilibria depends on the RNV, which is especially large ($\approx 0.3$) against the value of the equilibrium for $E_4$ (which is between 7.6 and 8.6) compared to other enzymes, causing this magnifying effect of the noise.

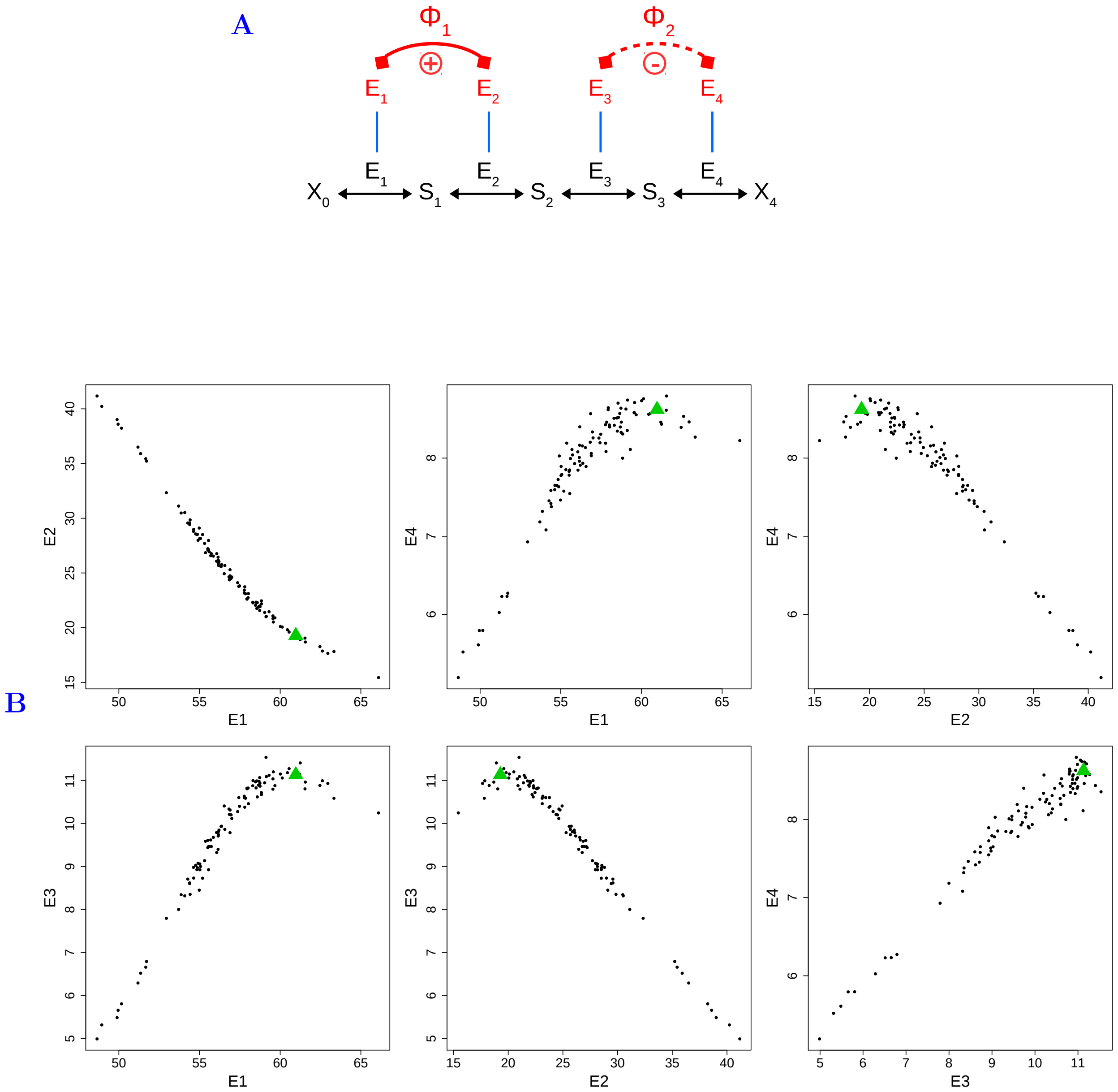

**Figure S4. End of simulation of enzyme evolution in a four-enzyme pathway when there are one positive and one negative groups with competition.** (A) The four-enzyme pathway used in these simulations. The enzymes $E_1$ and $E_2$ are positively co-regulated, whereas enzymes $E_3$ and $E_4$ are negatively co-regulated. Parameter values: $\mathbf{A} = (1, 10, 30, 50)$, $X = 1$, $N = 10000$, $\beta_{12} = 0.32$, $\beta_{43} = -0.43$. (B) Relationship between enzyme concentrations at the end of the simulations. One hundred simulations were performed, with different initial concentrations. Each black point correspond to the end of one simulation. The green triangles ($E_1 = 61$, $E_2 = 19.3$, $E_3 = 11.1$, $E_4 = 8.6$) correspond to the maximum flux without regulation. The black points seem to describe a curve, with noise that is likely due to different sizes of the neutral zones. Although we cannot estimate the equilibrium, it is totally deterministic and depends on the initial concentrations.

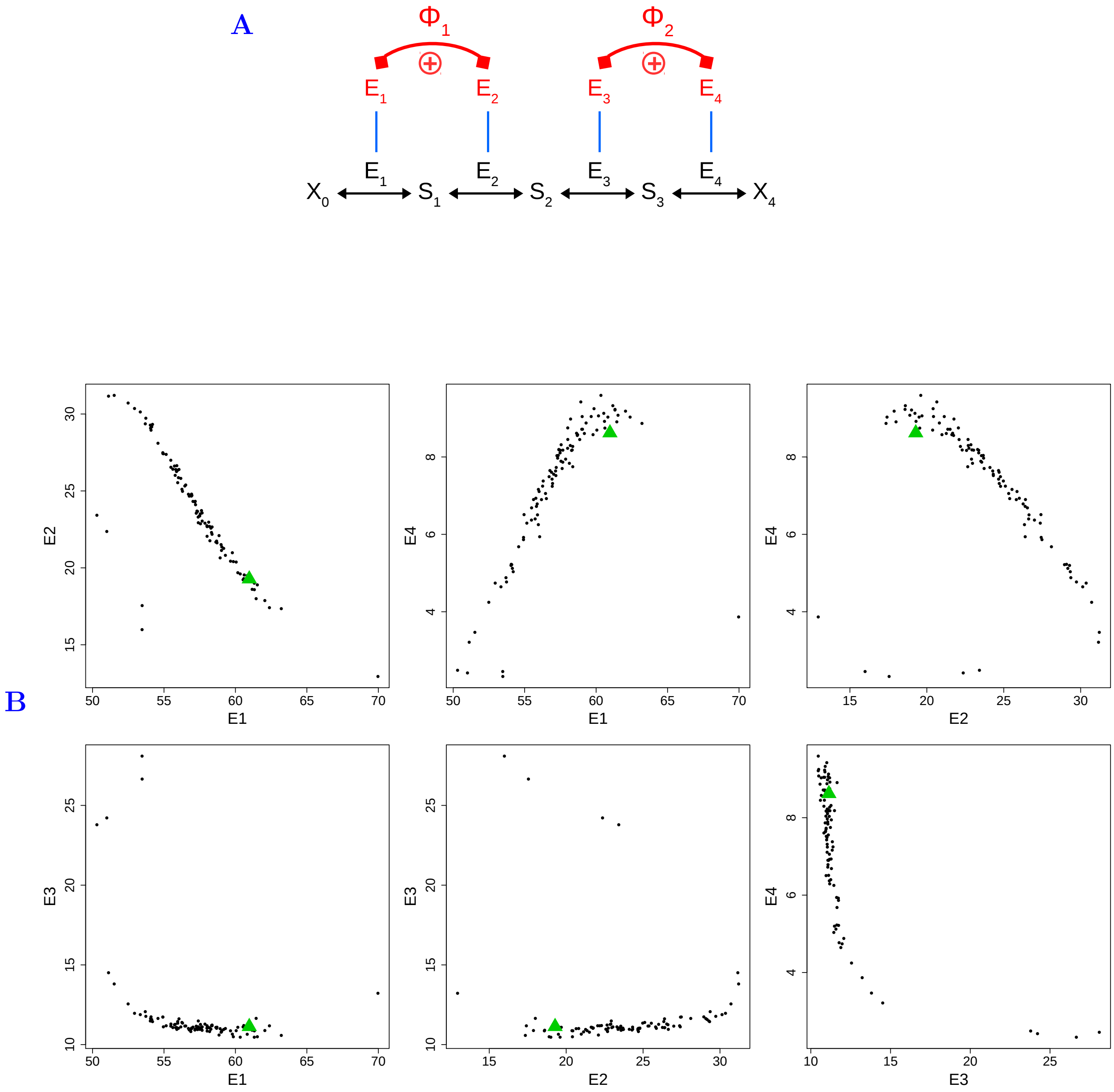

**Figure S5. End of simulation of enzyme evolution in a four-enzyme pathway when there are two positive groups with competition.** (A) The four-enzyme pathway used in these simulations. Enzymes $E_1$ and $E_2$ are positively co-regulated, as are enzymes $E_3$ and $E_4$. Parameter values: $\mathbf{A} = (1, 10, 30, 50)$, $X = 1$, $N = 10000$, $\beta_{12} = 0.32$, $\beta_{34} = 0.1$. (B) Relationship between the enzyme concentrations at the end of simulation. One hundred simulations were performed, with different initial concentrations. The symbols are the same as figure S4. As in figure S4, the black points seem to describe a curve, with noise that is likely due to different sizes of the neutral zones. Although we cannot estimate the equilibrium, it is totally deterministic and depends on the initial concentration.

### II Mathematical Proofs

#### II.1 Properties of co-regulation groups

$\Phi_q$ is the co-regulation group $q$, with $q \in \{1, p\}$, and $m_q$ is the number of enzymes in $\Phi_q$. The total number of enzymes is $n$, so $\sum_{q=1}^{n} m_q = n$. There is at least one enzyme by group and at most $n - p + 1$ enzymes, *i.e.* $\forall q, 1 \leq m_q \leq n - p + 1$.

If there is only one enzyme in a group ($m_q = 1$), this enzyme is independent from all other enzymes.

If there is only one group ($p = 1, m_1 = n$), all enzymes are in this group and they are co-regulated. This is the case "Co-regulation between all enzymes".

If all enzymes are independent ($p = n, \forall q\ m_q = 1$), there are as many groups as enzymes. This is the case "Independence between all enzymes".

#### II.2 Searching for evolutionary equilibrium

We search for the evolutionary equilibrium of total relative enzyme concentrations:

$$\forall j, \frac{\partial e_j}{\partial t} = 0$$

where $e_j = E_j/E_{\text{tot}}$. The total relative enzyme concentration can be partitioned into intra-group relative concentration $e_j^q = E_j/E^q$, where $E_q = \sum_{j \in \Phi_q} E_j$, and inter-group relative concentration $e_q = E_q/E_{\text{tot}}$.

Because $e_j = e_j^q e^q$, we have:

$$\frac{\partial e_j}{\partial t} = \frac{\partial (e_j^q e^q)}{\partial t} = \frac{\partial e_j^q}{\partial t} e^q + e_j^q \frac{\partial e^q}{\partial t} \tag{S1}$$

Thus, solving $\frac{\partial e_j}{\partial t} = 0$ is equivalent to solving $\frac{\partial e_j^q}{\partial t} = 0$ *and* $\frac{\partial e^q}{\partial t} = 0$.

$$\boxed{\frac{\partial e_j}{\partial t} = 0 \Longleftrightarrow \frac{\partial e_i^q}{\partial t} = 0 \quad \& \quad \frac{\partial e^q}{\partial t} = 0} \tag{S2}$$

#### II.3 Equilibria when there is no competition

Without competition for resources (free $E_{\text{tot}}$), the redistribution coefficient is $\alpha_{ij} = \beta_{ij}$ for all $(i, j)$, and the values of the redistribution coefficient are:

$$\forall q,\ \forall i \in \Phi_q,\ \forall j \quad \alpha_{ij} = \beta_{ij} = \begin{cases} 1/\beta_{ji} & \text{if } j \in \Phi_q \\ 1 & \text{if } j = i \\ 0 & \text{if } j \notin \Phi_q \end{cases} \tag{S3}$$

So the differential system (equation 7) that describes the evolution of enzyme concentrations becomes:

$$\frac{\partial E_j}{\partial t} = 2\mu \sum_{i \in \Phi_q} s_i \beta_{ij} \delta_i \tag{S4}$$

And the flux response coefficient writes for all $i$ in $\Phi_q$:

$$R^J_{E_i} = E_i \frac{\sum\limits_{j=1}^{n} \frac{\beta_{ij}}{A_j E_j^2}}{\sum\limits_{j=1}^{n} \frac{1}{A_j E_j}} = e_i \frac{\sum\limits_{j\in\Phi_q} \frac{\beta_{ij}}{A_j e_j^2}}{\sum\limits_{j=1}^{n} \frac{1}{A_j e_j}} \tag{S5}$$

##### II.3.1 Equilibrium within co-regulation groups

**II.3.1.1 Condition for equilibrium** We search $\forall q$, $\forall j \in \Phi_q$, $e_j^{q*}$ such as $\frac{\partial e_j^q}{\partial t} = 0$, with $e_j^q = E_j/E^q$. We have:

$$\frac{\partial e_j^q}{\partial t} = \frac{\partial (E_j/E^q)}{\partial t} = \frac{\frac{\partial E_j}{\partial t} E^q - E_j \frac{\partial E^q}{\partial t}}{(E^q)^2} \tag{S6}$$

From equation S4, we can write for any co-regulation group $\Phi_q$:

$$\begin{aligned}\frac{\partial E^q}{\partial t} = \frac{\partial \left( \sum\limits_{j\in\Phi_q} E_j \right)}{\partial t} &= \sum_{j\in\Phi_q} \frac{\partial E_j}{\partial t} \\ &= \sum_{j\in\Phi_q} 2\mu \sum_{i\in\Phi_q} s_i \beta_{ij} \delta_i \\ &= 2\mu \sum_{i\in\Phi_q} s_i \delta_i \sum_{j\in\Phi_q} \beta_{ij}\end{aligned}$$

As the global co-regulation coefficient is $B_i = \sum\limits_{j\in\Phi_q} \beta_{ij}$ for all $i \in \Phi_q$, we can write:

$$\frac{\partial E^q}{\partial t} = 2\mu \sum_{i\in\Phi_q} s_i \delta_i B_i \tag{S7}$$

By replacing derivatives with their expression in equation (S6), we obtain:

$$\begin{aligned}\frac{\partial e_j^q}{\partial t} &= \frac{E^q 2\mu \sum\limits_{i\in\Phi_q} s_i \beta_{ij} \delta_i - E_j 2\mu \sum\limits_{i\in\Phi_q} \delta_i s_i B_i}{(E^q)^2} \\ &= \frac{2\mu}{E^q} \left( \sum_{i\in\Phi_q} \beta_{ij} s_i \delta_i - e_j^q \sum_{i\in\Phi_q} s_i \delta_i B_i \right) \\ &= \frac{2\mu}{E^q} \sum_{i\in\Phi_q} s_i \delta_i (\beta_{ij} - e_j^q B_i)\end{aligned}$$

Using the relationship between the selection coefficient and flux response coefficient $s_i = R^J_{E_i} \delta_i / E_i$ (Coton et al. 2021), and the relationship $e_i^q = E_i/E^q$, we get:

$$\begin{aligned}\frac{\partial e_j^q}{\partial t} &= \frac{2\mu}{E^q} \sum_{i\in\Phi_q} R^J_{E_i} \frac{\delta_i^2}{E_i} (\beta_{ij} - e_j^q B_i) \\ &= \frac{2\mu}{(E^q)^2} \sum_{i\in\Phi_q} \frac{R^J_{E_i} \delta_i^2}{e_i^q} (\beta_{ij} - e_j^q B_i)\end{aligned}$$

and from the relationship $B_i = \beta_{ij} B_j$, which is valid within groups, we get:

$$\frac{\partial e_j^q}{\partial t} = \frac{2u}{(E^q)^2} \sum_{i \in \Phi_q} \frac{R_{E_i}^J \delta_i^2}{e_i^q} \left( \frac{B_i}{B_j} - e_j^q B_i \right)$$
$$= \frac{2u}{(E^q)^2} \left( \frac{1}{B_j} - e_j^q \right) \sum_{i \in \Phi_q} \frac{R_{E_i}^J \delta_i^2}{e_i^q} B_i$$

Therefore, the condition for evolutionary equilibrium within groups is:

$$\boxed{\frac{2u}{(E^q)^2} \left( \frac{1}{B_j} - e_j^q \right) \sum_{i \in \Phi_q} \frac{R_{E_i}^J \delta_i^2}{e_i^q} B_i = 0} \tag{S8}$$

**II.3.1.2 Theoretical equilibrium within groups** An obvious solution for equation S8 is:

$$\boxed{\forall i \in \Phi_q, \quad e_i^{q*} = \frac{1}{B_i}} \tag{S9}$$

Note that if at least one co-regulation coefficient is negative, leading to at least one negative $B_i$ in the group, the equilibrium concentration is negative, which is not realistic.

**II.3.1.3 Effective equilibrium within groups** Actually equation S8 has another trivial solution:

$$\boxed{\forall i \ \Phi_q \quad R_{E_i}^J = 0} \tag{S10}$$

or, from equation S5:

$$\forall i \in \Phi_q \quad \sum_{j \in \Phi_q}^{n} \frac{\beta_{ij}}{A_j (e_j)^2} = 0$$

As $e_j = e_j^q e^q$, we get:

$$\forall i \in \Phi_q \quad \frac{1}{(e^q)^2} \sum_{j \in \Phi_q}^{n} \frac{\beta_{ij}}{A_j (e_j^q)^2} = 0$$

This equality can only be solved if at least one co-regulation coefficient is negative in group $\Phi_q$. Because $\forall (i,j) \in \Phi_q, \beta_{ij} = B_i / B_j$ (equation 5), we get:

$$\forall i \in \Phi_q \quad B_i \sum_{j \in \Phi_q} \frac{1}{B_j A_j (e_j)^2} = 0$$

or

$$\boxed{\sum_{j \in \Phi_q} \frac{1}{B_j A_j (\tilde{e_j^q})^2} = 0} \tag{S11}$$

where $\tilde{e_j^q}$ is the value at intra-group effective equilibrium that solved this equation. Thanks to the relationship between $e_j^q$ and $\tau^q$ (equation 6), searching for $\tilde{e_j^q}$ for all $j \in \Phi_q$ is equivalent to searching for $\tilde{\tau}^q$ using the following equation:

$$\sum_{j \in \Phi_q} \frac{1}{B_j A_j (\tilde{\tau^q}(1/B_j - e_j^{q0}) + e_j^{q0})^2} = 0 \tag{S12}$$

##### II.3.2 Equilibrium between co-regulation groups

**II.3.2.1 Condition for equilibrium** We search $\forall q$, $e^{q*}$ such as $\frac{\partial e^q}{\partial t} = 0$, with $e^q = E^q/E_{\text{tot}}$. We have:

$$\frac{\partial e^q}{\partial t} = \frac{\partial(E^q/E_{tot})}{\partial t} = \frac{\frac{\partial E^q}{\partial t}E_{tot} - E^q\frac{\partial E_{tot}}{\partial t}}{(E_{tot})^2} \tag{S13}$$

From equation S4, we can write for any co-regulation group $\Phi_q$ (equation S7, Supporting Information II.3.1.1):

$$\frac{\partial E^q}{\partial t} = 2\mu \sum_{i\in\Phi_q} s_i\delta_i B_i$$

In a same way, we get for the total concentration:

$$\frac{\partial E_{\text{tot}}}{\partial t} = \sum_{j=1}^{n} \frac{\partial E_j}{\partial t} = \sum_{q=1}^{p} \frac{\partial E^q}{\partial t} = \sum_{q=1}^{p} 2\mu \sum_{i\in\Phi_q} s_i\delta_i B_i$$

or

$$\frac{\partial E_{\text{tot}}}{\partial t} = 2\mu \sum_{i=1}^{n} s_i\delta_i B_i$$

By replacing derivatives with their expression in equation (S13), we obtain:

$$\begin{aligned}\frac{\partial e^q}{\partial t} &= \frac{E_{tot}2\mu \sum\limits_{i\in\Phi_q} s_i\delta_i B_i - E^q 2\mu \sum\limits_{i=1}^{n} s_i\delta_i B_i}{(E_{tot})^2}\\ &= \frac{2\mu}{E_{\text{tot}}}\left(\sum_{i\in\Phi_q} s_i\delta_i B_i - e^q \sum_{i=1}^{n} s_i\delta_i B_i\right)\end{aligned}$$

Using $s_i = R^J_{E_i}\delta_i/E_i$ (Coton et al. 2021), we get:

$$\frac{\partial e^q}{\partial t} = \frac{2\mu}{E_{\text{tot}}}\left(\sum_{i\in\Phi_q} R^J_{E_i}\frac{\delta_i^2}{E_i}B_i - e^q \sum_{i=1}^{n} R^J_{E_i}\frac{\delta_i^2}{E_i}B_i\right)$$

When there is co-regulation without competition, we have $\delta_i = \nu$ for all $i$ (Coton et al. 2021), so:

$$\frac{\partial e^q}{\partial t} = \frac{2\mu\nu^2}{E_{\text{tot}}}\left(\sum_{i\in\Phi_q} \frac{R^J_{E_i}}{E_i}B_i - e^q \sum_{i=1}^{n} \frac{R^J_{E_i}}{E_i}B_i\right)$$

Thus, the condition $\dfrac{\partial e^q}{\partial t} = 0$ for evolutionary equilibrium between groups is:

$$\boxed{\frac{1}{e^q}\sum_{i\in\Phi_q} \frac{R^J_{E_i}}{E_i}B_i = \sum_{i=1}^{n} \frac{R^J_{E_i}}{E_i}B_i} \tag{S14}$$

**II.3.2.2 Positive groups and singletons** To make apparent the other co-regulation groups in equation S14, we split the sum of the right side of the equation:

$$\frac{1}{e^q}\sum_{i\in\Phi_q}\frac{R^J_{E_i}}{E_i}B_i=\sum_{k=1}^{p}\sum_{i\in\Phi_k}\frac{R^J_{E_i}}{E_i}B_i$$

Replacing the response coefficient by its expression (equation S5), we get:

$$\frac{1}{e^q}\sum_{i\in\Phi_q}B_i\sum_{j\in\Phi_q}\frac{\beta_{ij}}{A_jE_j^2}=\sum_{k=1}^{p}\sum_{i\in\Phi_k}B_i\sum_{j\in\Phi_k}\frac{\beta_{ij}}{A_jE_j^2}$$

Thus we can use the relationship $\beta_{ij}=B_i/B_j$ (equation 5), which is valid only if $i$ and $j$ are in the same co-regulation group:

$$\frac{1}{e^q}\sum_{i\in\Phi_q}B_i^2\sum_{j\in\Phi_q}\frac{1}{B_jA_jE_j^2}=\sum_{k=1}^{p}\sum_{i\in\Phi_k}B_i^2\sum_{j\in\Phi_k}\frac{1}{B_jA_jE_j^2}$$

From the relationships $e_j^q=E_j/E^q$ and $e^q=E^q/E_{\text{tot}}$, we get:

$$\frac{1}{e^q}\sum_{i\in\Phi_q}B_i^2\sum_{j\in\Phi_q}\frac{1}{B_jA_j(e_j^qe^qE_{\text{tot}})^2}=\sum_{k=1}^{p}\sum_{i\in\Phi_k}B_i^2\sum_{j\in\Phi_k}\frac{1}{B_jA_j(e_j^ke^kE_{\text{tot}})^2}$$

The inter-group relative concentrations $e^q$ and $e^k$ can be taken out from the sums:

$$\frac{1}{(e^q)^3}\sum_{i\in\Phi_q}B_i^2\sum_{j\in\Phi_q}\frac{1}{B_jA_j(e_j^q)^2}=\sum_{k=1}^{p}\frac{1}{(e^k)^2}\sum_{i\in\Phi_k}B_i^2\sum_{j\in\Phi_k}\frac{1}{B_jA_j(e_j^k)^2}$$

As we consider positive groups and singletons, the intra-group theoretical equilibrium is $e_j^{q*}=1/B_j$ (equation 11), so:

$$\frac{1}{(e^{q*})^3}\sum_{i\in\Phi_q}B_i^2\sum_{j\in\Phi_q}\frac{B_j}{A_j}=\sum_{k=1}^{p}\frac{1}{(e^{k*})^2}\sum_{i\in\Phi_k}B_i^2\sum_{j\in\Phi_k}\frac{B_j}{A_j}$$

Let's call $A^{q*}$ the *apparent activity of co-regulation group* $\Phi_q$ at theoretical equilibrium:

$$\boxed{A^{q*}=\left(\sum_{\substack{i\in\Phi_q\\ j\in\Phi_q}}\frac{B_i^2B_j}{A_j}\right)^{-1}} \tag{S15}$$

So the previous expression becomes:

$$\frac{1}{A^{q*}(e^{q*})^3}=\sum_{k=1}^{p}\frac{1}{A^{k*}(e^{k*})^2}$$

As $\sum_{k=1}^{n}\frac{1}{A^{k*}(e^{k*})^2}$ is constant, this equality is valid for any co-regulation group:

$$\frac{1}{A^{q*}(e^{q*})^3}=\frac{1}{A^{q'*}(e^{q'*})^3}=\sum_{k=1}^{p}\frac{1}{A^{k*}(e^{k*})^2}$$

Because the apparent activities are strictly positive when there are only positive groups and singletons, we get:

$$\forall q,\ \forall q', \quad e^{q'*} = e^{q*}\frac{(A^q)^{1/3}}{(A^{q'})^{1/3}}$$

Since $\sum\limits_{q'=1}^{p} e^{q'} = 1$, we have:

$$\sum_{q'=1}^{p} e^{q'*} = e^{q*}(A^{q*})^{1/3}\sum_{q'=1}^{p}\frac{1}{(A^{q'*})^{1/3}} = 1$$

This leads to the expression of the theoretical equilibrium of relative enzyme concentrations when all groups are positives or singletons:

$$\boxed{\forall q \quad e^{q*} = \frac{(1/A^{q*})^{1/3}}{\sum\limits_{q'=1}^{p}(1/A^{q'*})^{1/3}}} \tag{S16}$$

**II.3.2.3 Negative groups** When all groups are negative, the flux response coefficients in equation S14 are all null, so we could not use the previous method to find the equilibria. To overcome the problem, we relied on the relationship between $\tau^q$ and the absolute concentrations within groups, $E^q$ (Supporting Information II.3.3. Thus, at effective equilibrium, the absolute concentration writes:

$$\tilde{E^q} = \frac{E^{q0}}{1-\tilde{\tau^q}} \tag{S17}$$

where $\tilde{\tau^q}$ is the value of the driving variable at the effective equilibrium. Therefore, the inter-group relative concentrations at the effective equilibrium are:

$$\boxed{\forall q \quad \tilde{e^q} = \frac{\frac{E^{q0}}{1-\tilde{\tau^q}}}{\sum\limits_{k=1}^{p}\frac{E^{k0}}{1-\tilde{\tau^k}}}} \tag{S18}$$

**II.3.2.4 Positive and negative groups** When there are both types of groups, the response coefficients are null for negative groups and non-null for positive groups and singletons. So equation S14 is solvable only for positive groups and singletons, which leads to this equilibrium between these groups:

$$\forall q \text{ such as } \theta_q \geq 0 \quad \frac{E^q}{\sum\limits_{\substack{q'=1\\ \theta_{q'}\geq 0}}^{p} E^{q'}} = \frac{(1/A^{q*})^{1/3}}{\sum\limits_{\substack{q'=1\\ \theta_{q'}\geq 0}}^{p}(1/A^{q'*})^{1/3}} \tag{S19}$$

which is similar to equation 12, by considering $E^q/\sum\limits_{\theta_q\geq 0} E^q$ instead of $e^q$.

In the negative groups the relative enzyme concentrations tend to their effective equilibrium.

To study the behavior of absolute concentrations, we use the relationship between absolute concentration and the driving variable (Supporting Information II.3.3):

$$\forall i \in \Phi_q \quad E_i = E_i^0 + \frac{\tau^q E^{q^0}}{(1-\tau^q)B_i} \tag{S20}$$

In the positive groups, $e_i^q$ tends to the intra-group theoretical equilibrium $1/B_i$, so $\tau^q$ tends to 1 (see equation 6), therefore $E_i$ tends to infinity. The enzyme concentration in singletons also increase to infinity (Coton et al. 2021). In the negative groups, $e_i^q$ reaches the effective equilibrium, so $\tau^q$ tends to $\tilde{\tau^q}$, therefore $E_i$ tends to $\tilde{E_i}$. Therefore, the absolute concentrations of enzymes in the negative groups become negligible compared to those in the positive groups and singletons. The inter-group relative concentrations tend towards the equilibrium of expression S19 in positive groups and towards zero in negative groups. Because $\sum_{\theta_q \geq 0} E^q$ tends towards $E_{\text{tot}}$, we can say that for positive groups and singletons $e^q$ tend towards $e^{q*}$ (equation 12) considering only positive groups and singletons in denominator instead of all groups, *i.e.*:

$$e^{q*} = \frac{(1/A^{q*})^{1/3}}{\sum\limits_{\substack{q'=1 \\ \theta_{q'} \geq 0}}^{p} (1/A^{q'*})^{1/3}} \tag{S21}$$

The same applies for $e_i$ that tend towards $e_i^*$ (equation 14).

When there are positive and negative co-regulation groups, the flux writes:

$$J = \frac{X}{\sum\limits_{j=1}^{n} \frac{1}{A_j E_j}} \tag{S22}$$

$$= \frac{X}{\sum\limits_{q=1}^{p} \sum\limits_{j \in \Phi_q}^{n} \frac{1}{A_j E_j}} \tag{S23}$$

$$= \frac{X}{\sum\limits_{\substack{q=1 \\ \theta_q \geq 0}}^{p} \sum\limits_{j \in \Phi_q}^{n} \frac{1}{A_j E_j} + \sum\limits_{\substack{q=1 \\ \theta_q < 0}}^{p} \sum\limits_{j \in \Phi_q}^{n} \frac{1}{A_j E_j}} \tag{S24}$$

As enzyme concentrations are stuck at the effective equilibrium in negative groups while they increase indefinitely in positive groups and singletons, we have:

$$\boxed{J \xrightarrow[t \to \infty]{} \frac{X}{\sum\limits_{\substack{q=1 \\ \theta_q < 0}}^{p} \sum\limits_{\substack{j=1 \\ j \in \Phi_q}}^{n} \frac{1}{A_j \tilde{E}_j}} = \tilde{J}_{\max}} \tag{S25}$$

##### II.3.3 Relationship between absolute concentrations and driving variable

The intra-group relative concentrations in a group are linearly related. The parametric equation of the line $\mathcal{E}^q$ writes:

$$\forall i \in \Phi_q \quad e_i^q = \tau^q (1/B_i - e_i^{q^0}) - e_i^{q^0} \tag{S26}$$

where $\tau^q$ is the driving variable.

The absolute concentrations can also be expressed depending on $\tau^q$.

The relation between $E^q$ and $\tau^q$ can be derived from the relation $B_i = \dfrac{\Delta E^q}{\Delta E_i}$ (equation 4). We have:

$$\begin{aligned} E^q &= E^{q^0} + B_i(E_i - E_i^0) \\ &= E^q B_i e_i^q + E^{q^0}(1 - B_i e_i q^0) \\ &= E^{q^0} \frac{1 - B_i e_i^{q^0}}{1 - B_i e_i^q} \end{aligned}$$

Replacing $e_i^q$ by its expression depending on $\tau^q$ (equation 6), we get:

$$E^q = E^{q^0} \frac{1 - B_i e_i^{q^0}}{1 - B_i(\tau^q(1/B_i - e_i^{q^0}) + e_i^{q^0})}$$

which is simplified to (for $\tau^q \neq 1$):

$$E^q = \frac{E^{q^0}}{1 - \tau^q} \tag{S27}$$

$$\boxed{E^q = \frac{E^{q0}}{1 - \tau^q}} \tag{S28}$$

In the parametric equation 6, we can make visible the *absolute* enzyme concentrations, using the equation $e_i^q = E_i / E^q$:

$$\forall i, \; \frac{E_i}{E^q} = \tau \left( \frac{1}{B_i} - \frac{E_i^0}{E^{q^0}} \right) + \frac{E_i^0}{E^{q0}} \tag{S29}$$

Introducing in this equation the expression of $E^q$ (equation S28), we get:

$$\forall i \Phi_q \; \frac{E_i(1 - \tau^q)}{E^{q^0}} = \tau^q \left( \frac{1}{B_i} - \frac{E_i^0}{E^{q^0}} \right) + \frac{E_i^0}{E^{q^0}}$$

Thus:

$$E_i = E_i^0 + \frac{\tau^q E^{q^0}}{(1 - \tau^q) B_i}$$

Or:

$$\boxed{\forall i \in \Phi_q \quad E_i = E_i^0 + \frac{E^{q0}}{B_i} \left( -1 + \frac{1}{1 - \tau^q} \right)} \tag{S30}$$

#### II.4 Equilibria when there is competition

With competition for resources between all enzymes (fixed $E_{\text{tot}}$), we have for the co-regulation coefficients:

$$\forall q, \; \forall i \in \Phi_q, \; \forall j \quad \beta_{ij} = \begin{cases} 1/\beta_{ji} & \text{if } j \in \Phi_q \\ 1 & \text{if } j = i \\ 0 & \text{if } j \notin \Phi_q \end{cases} \tag{S31}$$

and for the redistribution coefficients:

$$\forall q,\ \forall i \in \Phi_q,\ \forall j \quad \alpha_{ij} = \frac{\beta_{ij}/B_i - e_j}{1/B_i - e_i} = \begin{cases} \dfrac{1/B_j - e_j}{1/B_i - e_i} & \text{if } j \in \Phi_q \\ \dfrac{-e_j}{1/B_i - e_i} & \text{if } j \notin \Phi_q \end{cases} \tag{S32}$$

So the system of differential equations that describes the evolution of enzyme concentrations (equation 7) becomes:

$$\begin{aligned} \forall q,\ \forall j \in \Phi_q \quad \frac{\partial E_j}{\partial t} &= 2\mu \sum_{i=1}^{n} s_i \delta_i \alpha_{ij} \\ &= 2\mu \sum_{i \in \Phi_q} s_i \delta_i \frac{1/B_j - e_j}{1/B_i - e_i} + 2\mu \sum_{i \notin \Phi_q} s_i \delta_i \frac{-e_j}{1/B_i - e_i} \end{aligned}$$

Thus:

$$\forall q,\ \forall j \in \Phi_q \quad \frac{\partial E_j}{\partial t} = 2\mu \frac{1}{B_j} \sum_{i \in \Phi_q} \frac{s_i \delta_i}{1/B_i - e_i} - 2\mu e_j \sum_{i=1}^{n} \frac{s_i \delta_i}{1/B_i - e_i} \tag{S33}$$

As the total concentration is fixed with competition, we have for the total relative concentration:

$$\forall q,\ \forall j \in \Phi_q \quad \frac{\partial e_j}{\partial t} = \frac{1}{E_{\text{tot}}} \frac{\partial E_j}{\partial t}$$

Thanks to the relationship between the selection coefficient and the response coefficient $s_i = R^J_{E_i} \delta_i / E_i$ (Coton et al. 2021), we get:

$$\forall q,\ \forall j \in \Phi_q \quad \frac{\partial e_j}{\partial t} = 2 \frac{\mu}{E_{\text{tot}}} \left( \frac{1}{B_j} \sum_{i \in \Phi_q} \frac{R^J_{E_i} \delta_i^2}{E_i (1/B_i - e_i)} - e_j \sum_{i=1}^{n} \frac{R^J_{E_i} \delta_i^2}{E_i (1/B_i - e_i)} \right) \tag{S34}$$

###### II.4.1 Effective equilibria

A solution to the evolutionary equilibrium $\dfrac{\partial e_j}{\partial t} = 0$ is:

$$\boxed{\forall i \quad R^J_{E_i} = 0} \tag{S35}$$

which is the definition of the effective equilibrium, and corresponds to maximal flux in regard to each enzyme.

$$\forall q,\ \forall i \in \Phi_q \quad R^J_{E_i} = 0 \Leftrightarrow e_i \frac{\sum_{j=1}^{n} \frac{\alpha_{ij}}{A_j e_j^2}}{\sum_{j=1}^{n} \frac{1}{A_j E_j}} = 0$$
$$\Leftrightarrow \sum_{j=1}^{n} \frac{\alpha_{ij}}{A_j e_j^2} = 0$$
$$\Leftrightarrow \sum_{j=1}^{n} \frac{\beta_{ij}/B_i - e_j}{1/B_i - e_i} \frac{1}{A_j e_j^2} = 0$$
$$\Leftrightarrow \sum_{j\in\Phi_q} \frac{1/B_j - e_j}{A_j e_j^2} + \sum_{j\notin\Phi_q} \frac{-e_j}{A_j e_j^2} = 0$$
$$\Leftrightarrow \sum_{j\in\Phi_q} \frac{1}{B_j A_j e_j^2} - \sum_{j=1}^{n} \frac{e_j}{A_j e_j^2} = 0$$

So the condition for the effective equilibrium is:

$$\forall q, \quad \sum_{j\in\Phi_q} \frac{1}{B_j A_j e_j^2} = \sum_{j=1}^{n} \frac{1}{A_j e_j} \tag{S36}$$

**II.4.1.1 Between groups** Replace $e_j$ by $e^q e^q_j$ in the left-hand side of equation S36:

$$\sum_{j\in\Phi_q} \frac{1}{B_j A_j e_j^2} = \sum_{j=1}^{n} \frac{1}{A_j e_j}$$
$$\Leftrightarrow \sum_{j\in\Phi_q} \frac{1}{B_j A_j (e^q e^q_j)^2} = \sum_{j=1}^{n} \frac{1}{A_j e_j}$$
$$\Leftrightarrow \frac{1}{(e^q)^2} \sum_{j\in\Phi_q} \frac{1}{B_j A_j (e^q_j)^2} = \sum_{j=1}^{n} \frac{1}{A_j e_j}$$

Defining $A^q$ as the apparent activity of co-regulation group $\Phi_q$, such as:

$$\frac{1}{A^q} = \sum_{j\in\Phi_q} \frac{1}{B_j A_j (e^q_j)^2} \tag{S37}$$

we get:

$$\frac{1}{(e^q)^2} \frac{1}{A^q} = \sum_{j=1}^{n} \frac{1}{A_j e_j} \tag{S38}$$

140 As $\sum_{j=1}^{n} \frac{1}{A_j e_j}$ is constant, this equality is valid for any group:

$$\frac{1}{(e^q)^2} \frac{1}{A^q} = \sum_{j=1}^{n} \frac{1}{A_j e_j} = \frac{1}{(e^k)^2} \frac{1}{A^k}$$

and:

$$(e^q)^2 = \frac{A^k (e^k)^2}{A^q}$$

Assuming that for all $q$ $A^q$ is positive even for negative groups (which has been empirically verify with some simulations when equilibrium is computable), we have:

$$e^k = \left( e^q \frac{A^q}{A^k} \right)^{1/2}$$

As $\sum_{k=1}^{p} e^k = 1$, we can write:

$$\sum_{k=1}^{p} e^k = 1 = \sum_{k=1}^{p} e^q \left( \frac{A^q}{A^k} \right)^{1/2}$$

So at the effective equilibrium, we have:

$$\boxed{\tilde{e^q} = \frac{\tilde{A^q}^{-1/2}}{\sum_{k=1}^{p} \tilde{A^k}^{-1/2}}} \tag{S39}$$

with

$$\frac{1}{\tilde{A^q}} = \sum_{j \in \Phi_q} \frac{1}{B_j A_j (\tilde{e^q_j})^2} \tag{S40}$$

###### II.4.1.2 Within groups

In expression S36, we replace $e_j$ by $e^q e^q_j$:

$$\frac{1}{(e^q)^2} \sum_{j \in \Phi_q} \frac{1}{B_j A_j (e^q_j)^2} - \sum_{k=1}^{p} \frac{1}{e^k} \sum_{j \in \Phi_k} \frac{1}{A_j e^k_j} = 0$$

In the first member, we recognize the expression of the apparent activity (equation S37), so we have:

$$\frac{1}{A^q (e^q)^2} - \sum_{k=1}^{p} \frac{1}{e^k} \sum_{j \in \Phi_k} \frac{1}{A_j e^k_j} = 0$$

Replacing $e^k$ by its expression (equation S39), we get:

$$\frac{1}{A^q \left( \frac{(A^q)^{-1/2}}{\sum_{q'=1}^{p} (\tilde{A^{q'}})^{-1/2}} \right)^2} - \sum_{k=1}^{p} \frac{1}{\frac{(A^k)^{-1/2}}{\sum_{q'=1}^{p} (\tilde{A^{q'}})^{-1/2}}} \sum_{j \in \Phi_k} \frac{1}{A_j e^k_j} = 0$$

Leading to:

$$\sum_{q'=1}^{p} (\tilde{A^{q'}})^{-1/2} - \sum_{k=1}^{p} (A^k)^{1/2} \sum_{j \in \Phi_k} \frac{1}{A_j e^k_j} = 0$$

However, if there is only one enzyme in $\Phi_q$, *i.e.* if $\theta_q = 0$, we have $e_i^q = 1$ and $A^q = A_i$. Thus $A_i^{-1/2} - A_i^{1/2}/A_i = A_i^{-1/2} - A_i^{-1/2} = 0$, and the elements corresponding to $\theta_q = 0$ disappear in the expression:

$$\boxed{\sum_{\substack{k=1\\ \theta_k\neq 0}}^{p} \left( (A^k)^{-1/2} - (A^k)^{1/2} \sum_{j\in\Phi_k} \frac{1}{A_j e_j^k} \right) = 0} \tag{S41}$$

**II.4.1.3 Particular case: only one regulated group and singletons** When there is only one co-regulated group $\Phi_q$ (positive or negative) and independent enzymes, we get from equation S41:

$$(A^q)^{-1/2} - (A^q)^{1/2} \sum_{j\in\Phi_q} \frac{1}{A_j e_j^q} = 0$$

or:

$$\frac{1}{A^q} = \sum_{j\in\Phi_q} \frac{1}{A_j e_j^q}$$

Using the expression of $A^q$ (equation S37), we get:

$$\sum_{j\in\Phi_q} \frac{1}{B_j A_j (e_j^q)^2} = \sum_{j\in\Phi_q} \frac{1}{A_j e_j^q}$$

$$\Leftrightarrow \sum_{j\in\Phi_q} \left( \frac{1}{B_j A_j (e_j^q)^2} - \frac{1}{A_j e_j^q} \right) = 0$$

$$\Leftrightarrow \sum_{j\in\Phi_q} \left( \frac{1/B_j - e_j^q}{A_j (e_j^q)^2} \right) = 0$$

Replacing $e_j^q$ by its expression in the parametric equation of the line $\mathcal{E}^q$ (equation 6), we get:

$$\sum_{j\in\Phi_q} \left( \frac{1/B_j - (\tau^q(1/B_j - e_j^{q^0}) - e_j^{q^0})}{A_j (e_j^q)^2} \right) = 0$$

$$\Leftrightarrow (1-\tau^q) \sum_{j\in\Phi_q} \left( \frac{1/B_j - e_j^{q^0}}{A_j (e_j^q)^2} \right) = 0$$

$$\boxed{(1-\tau^q) \sum_{j\in\Phi_q} \frac{1/B_j - e_j^{q0}}{A_j(\tau^q(1/B_j - e_j^{q0}) + e_j^{q0})^2} = 0} \tag{S42}$$

Thus, there are two solutions:

- $\tau^{q*} = 1$, which is the theoretical equilibrium (Supporting Information II.4.2), but in this case the flux response coefficient is not null
- $\tilde{\tau^q} \neq 1$, which cancels the right part of the equation and can be found numerically. In this case, $\tilde{\tau^q}$ nullifies the response coefficient, which corresponds to a local maximal flux. The right part of the equation is similar to the expression of the effective equilibrium when all enzymes are co-regulated.

##### II.4.2 Intra-group theoretical equilibria

Here we search $e_i^{q*}$ such as $\dfrac{\partial e_i^q}{\partial t} = 0$.

$$\forall q,\ \forall j \in \Phi_q, \quad \frac{\partial e_j^q}{\partial t} = \frac{\partial E_j / E^q}{\partial t} = \frac{E^q \dfrac{\partial E_j}{\partial t} - E_j \dfrac{\partial E^q}{\partial t}}{(E^q)^2} \tag{S43}$$

From this equation S33, we can write for any co-regulation group $\Phi_q$:

$$\begin{aligned}
\forall q, \quad \frac{\partial E^q}{\partial t} &= \frac{\partial \left( \sum\limits_{j\in\Phi_q} E_j \right)}{\partial t} = \sum_{j\in\Phi_q} \frac{\partial E_j}{\partial t} \\
&= \sum_{j\in\Phi_q} \left( 2\mu \frac{1}{B_j} \sum_{i\in\Phi_q} \frac{s_i\delta_i}{1/B_i - e_i} - 2\mu e_j \sum_{i=1}^{n} \frac{s_i\delta_i}{1/B_i - e_i} \right) \\
&= 2\mu \left( \sum_{i\in\Phi_q} \frac{s_i\delta_i}{1/B_i - e_i} \sum_{j\in\Phi_q} \frac{1}{B_j} - \sum_{i=1}^{n} \frac{s_i\delta_i}{1/B_i - e_i} \sum_{j\in\Phi_q} e_j \right)
\end{aligned}$$

As $\sum\limits_{j\in\Phi_q} e_j = \sum\limits_{j\in\Phi_q} \dfrac{E_j}{E_{\text{tot}}} = \dfrac{E^q}{E_{\text{tot}}} = e^q$ and $\sum\limits_{j\in\Phi_q} \dfrac{1}{B_j} = 1$, we get:

$$\forall q, \quad \frac{\partial E^q}{\partial t} = 2\mu \left( \sum_{i\in\Phi_q} \frac{s_i\delta_i}{1/B_i - e_i} - e^q \sum_{i=1}^{n} \frac{s_i\delta_i}{1/B_i - e_i} \right) \tag{S44}$$

Replacing $\dfrac{\partial E_j}{\partial t}$ and $\dfrac{\partial E^q}{\partial t}$ by their expressions (respectively S33 and S44) in equation S43, and with $e_j^q = E_j/E^q$, we have:

$$\frac{\partial e_j^q}{\partial t} = \frac{2\mu}{E^q} \left[ \left( \frac{1}{B_j} \sum_{i\in\Phi_q} \frac{s_i\delta_i}{1/B_i - e_i} - e_j \sum_{i=1}^{n} \frac{s_i\delta_i}{1/B_i - e_i} \right) - e_j^q \left( \sum_{i\in\Phi_q} \frac{s_i\delta_i}{1/B_i - e_i} - e^q \sum_{i=1}^{n} \frac{s_i\delta_i}{1/B_i - e_i} \right) \right]$$

Because $e_i = e_i^q e^q$, we get:

$$\frac{\partial e_j^q}{\partial t} = \frac{2\mu}{E^q} \left[ \left( \frac{1}{B_j} - e_j^q \right) \sum_{i\in\Phi_q} \frac{s_i\delta_i}{1/B_i - e_i} \right]$$

There is an obvious solution for $\dfrac{\partial e_j^q}{\partial t} = 0$, which is the theoretical equilibrium within groups:

$$\boxed{\forall j \in \Phi_q, \quad e_j^{q*} = \frac{1}{B_j}} \tag{S45}$$

This expression of $e_i^q$ solves $\partial e_i^q/\partial t = 0$, but does not solve $\partial e_i/\partial t = 0$. However, if at least one co-regulation coefficient is negative, leading to at least one negative $B_i < 0$ in the group, the equilibrium concentration is negative, which is impossible to reach.

#### II.5 Range of neutral variation without competition

To search if the selection coefficient tends to zero at equilibrium, which corresponds to neutrality of variations of enzyme concentrations, we used the relationship between the flux response coefficients and the selection coefficients (Coton et al. 2021):

$$s_i = R^J_{E_i} \frac{\delta_i}{E_i} \tag{S46}$$

##### II.5.1 When there are only positive groups and singletons

At the theoretical equilibrium, the response coefficients are positive (using 14 in equation S5), and $\beta_{ij} \geq 0$ for all $(i, j)$), whereas $E_i$ can increase indefinitely. Therefore the selection coefficients tends to zero.

##### II.5.2 When there are only negative groups

At the effective equilibrium the response coefficients are null for all enzymes, therefore the selection coefficients are null.

##### II.5.3 When there are positive (and/or singletons) and negative groups

When there are both types of groups, the response coefficients depend on the group in which are the enzymes, according to the relation (equation S5):

$$\forall i \in \Phi_q, \quad R^J_{E_i} = e_i \frac{\sum\limits_{j \in \Phi_q} \frac{\beta_{ij}}{A_j e_j^2}}{\sum\limits_{j=1}^{n} \frac{1}{A_j e_j}}$$

At equilibrium for positive groups and singletons, the numerator is strictly positive and tends toward a constant if $\theta_q \geq 0$. For negative groups, it is null if $\theta_q < 0$.

The denominator is the same for all enzymes, whatever their co-regulation groups. It can be expanded as follows:

$$\begin{aligned}
\sum_{j=1}^{n} \frac{1}{A_j e_j} &= \sum_{q=1}^{p} \sum_{j \in \Phi_q} \frac{1}{A_j e_j^q e^q} \\
&= \sum_{\substack{q=1 \\ \theta_q \geq 0}}^{p} \frac{1}{e^q} \sum_{j \in \Phi_q} \frac{1}{A_j e_j^q} + \sum_{\substack{q=1 \\ \theta_q < 0}}^{p} \frac{1}{e^q} \sum_{j \in \Phi_q} \frac{1}{A_j e_j^q}
\end{aligned}$$

Replacing the $e_j^q$ by their respective equilibrium (equation 11 for $\theta_q \geq 0$ and equation 16 for $\theta_q < 0$), we get:

$$\sum_{j=1}^{n} \frac{1}{A_j e_j} = \sum_{\substack{q=1 \\ \theta_q \geq 0}}^{p} \frac{1}{e^q} \sum_{\substack{j=1 \\ E_j \in \Phi_q}}^{n} \frac{B_j}{A_j} + \sum_{\substack{q=1 \\ \theta_q < 0}}^{p} \frac{1}{e^q} \sum_{\substack{j=1 \\ E_j \in \Phi_q}}^{n} \frac{1}{A_j \widetilde{e_j^q}}$$

When the system tends to equilibrium within groups, (see section 3.1.3), $e^q \xrightarrow[t\to\infty]{} 0$ for $\theta_q < 0$ and $e^q \xrightarrow[t\to\infty]{} e^{q*}$ for $\theta_q \geq 0$. Thus, we have for the denominator:

$$\sum_{j=1}^{n} \frac{1}{A_j e_j} \xrightarrow[t\to\infty]{} +\infty$$

Therefore, we have for positive groups and singletons:

$$\forall i \in \Phi_q,\ \theta_q \geq 0, \quad R^J_{E_i} \xrightarrow[t\to\infty]{} e_i^* \frac{\sum\limits_{\substack{j=1 \\ E_j\in\Phi_q}}^{n} \frac{\beta_{ij}}{A_j (e_j^*)^2}}{+\infty} = 0^+$$

$$\begin{aligned} E_i &\xrightarrow[t\to\infty]{} \infty \\ \text{and} \quad R^J_{E_i} &\xrightarrow[t\to\infty]{} 0^+ \\ \text{so} \quad s_i \approx R^J_{E_i} \frac{\delta_i}{E_i} &\xrightarrow[t\to\infty]{} 0 \end{aligned}$$

For negative groups, the system tends toward effective equilibrium:

$$\begin{aligned} E_i &\xrightarrow[t\to\infty]{} \tilde{E}_i \\ \text{and} \quad R^J_{E_i} &\xrightarrow[t\to\infty]{} 0 \\ \text{so } s_i \approx R^J_{E_i} \frac{\delta_i}{E_i} &\xrightarrow[t\to\infty]{} 0 \end{aligned}$$

In conclusion, the selection coefficients tend to zero whatever the co-regulation groups.

##### II.5.4 RNV order

Combining equations S5 and S46 and because $\delta_i = \mu$ for all $i$ when there is no competition, the selection coefficients writes:

$$\forall i \in \Phi_q, \quad s_i \approx \frac{\mu}{E_i} E_i \frac{\sum\limits_{j\in\Phi_q} \frac{\beta_{ij}}{A_j E_j^2}}{\sum\limits_{j=1}^{n} \frac{1}{A_j E_j}}$$

The denominator is common for all enzymes, thus we write $Den = \sum\limits_{j=1}^{n} \frac{1}{A_j E_j}$. We use $E_i = E_{\text{tot}} e_i^q e^q$.

For positive groups and singletons, we have at theoretical equilibrium:

$$\forall i \in \Phi_q, \quad s_i \approx \mu \frac{\sum_{j\in\Phi_q} \frac{\beta_{ij}}{A_j E_j^2}}{Den}$$
$$\approx \frac{\mu}{E_{\text{tot}}^2 Den} \sum_{j\in\Phi_q} \frac{B_i/B_j}{A_j \left(\frac{1}{B_j}\right)^2 \left(\frac{(A^{q*})^{-1/3}}{\sum_{k=1}^{p}(A^{k*})^{-1/3}}\right)^2}$$
$$\approx \frac{\mu(\sum_{k=1}^{p}(A^{k*})^{-1/3})^2}{E_{\text{tot}}^2 Den} \sum_{j\in\Phi_q} \frac{B_i (A^{q*})^{2/3}}{A_j B_j}$$

The sum on the elements on the group is too complex to compare between groups. However, for independent enzymes, we have $B_i = 1$ and $A^q = A_j$, thus the selection coefficient becomes for singletons:

$$s_i \approx \frac{\mu(\sum_{k=1}^{p}(A^{k*})^{-1/3})^2}{E_{\text{tot}}^2 Den} A_j^{-1/3}$$

As RNV order is the reverse of selection coefficient order (Coton et al. 2021), we suggest that the RNV is in the same order as pseudo-activities between independent enzymes at theoretical equilibrium.

As in Coton et al. (2021), we consider the inferior and the superior limits of the RNV for enzyme $i$ in $\Phi_q$. Using the relationship between $E_i$ and $\tau^q$ (equation S30), we have:

$$|E_i^{\text{sup}} - E_i^{\text{sup}}| = |\frac{E^{q0}}{B_i} \left(\frac{1}{1-\tau^{q,textsup}} - \frac{1}{1-\tau^{q,\text{inf}}}\right)$$

where where $\tau^{q,textsup}$ and $\tau^{q,textinf}$ are the superior and inferior limits of the driving variable for group $\Phi_q$. Thus the RNVs of enzymes in the same group are inversely related to $|B_i|$'s.

However, we cannot compare the RNV between groups.

### III Complementary analysis

#### III.1 Degrees of freedom depend on the applied constraint

We showed that the evolutionary equilibrium and the flux shape depend on the constraint applied on the pathway.

In other words, each constraint modifies the adaptive landscape, by limiting the ability to move in the multidimensional space $(\mathbf{E}, J)$. We searched the number of degrees of freedom (df) of the system, which is the number of independent variables necessary and sufficient to describe the system, given the fixed parameters – here the pseudo-activities $\mathbf{A}$, the co-regulation coefficients of matrix $M_\beta$ and the initial concentrations $\mathbf{E^0}$. The number of df is fully determined by the relationship between enzyme concentrations, as the flux is a function of enzyme parameters (equation 1). Lower df means that the system lose dimensions on which he can move in the space $(\mathbf{E}, J)$, modifying the adaptive landscape. The diminution of df number depends on the applied constraint:

(i) In a system without any constraint on enzymes, there are as many degrees of freedom as enzymes. The system moves freely in the multidimensional space $(\mathbf{E}, J)$, although there is an optimal proportions of *relative* enzyme concentrations – the evolutionary equilibrium.

(ii) When there is competition between all enzymes, we only have to determine $n - 1$ enzyme concentrations to fully describe the system, because the total concentration is constant. The adaptive landscape becomes a dome in $(\mathbf{E}, J)$, the shape of which depending only on pseudo-activities (Coton et al. 2021).

(iii) When there is regulation between *all* enzymes, the enzyme concentrations are all linearly related, and fully determined by the driving variable $\tau$ (Coton et al. 2021): the system has one degree of freedom. Thus the adaptive landscape is reduced to two dimensions, $\tau$ and $J$. The landscape shape depends on the kind of co-regulation: flux increases indefinitely if enzymes are positively co-regulated, or reaches a maximum in case or negative co-regulation.

(iv) When there is competition plus co-regulation between *all* enzymes, the driving variable also fully describes the system, which so has one degree of freedom, and the adaptive landscape has two dimensions. However, the flux reaches a maximum whatever the kind of co-regulations, and the adaptive landscape is a section, oriented by the global co-regulation coefficients and the initial concentrations, of the dome of competition (Coton et al. 2021). Note that if the relative concentrations are initially equal to $1/\mathbf{B}$ ($B_i$ quantifies the effect on the total concentration of a variation of $E_i$ before applying the constraint on $E_{\text{tot}}$), the system is stuck at the theoretical equilibrium – an evolutionary black-hole, because at this point, the effect of a mutation is erased (see Coton et al. 2021). In this particular case, the system has zero degree of freedom and does not go outside this point.

(v) Here we worked on co-regulation groups. When there are co-regulation groups without competition, enzyme concentrations are linearly related within groups, and are independent from enzymes in other groups. So each co-regulation group can fully be described by its driving variable $\tau_q$. Note that for singleton groups, as the enzyme is independent from the others, its concentration replaces the driving variable and so represents a degree of freedom. Thus in this case, there is as many degrees of freedom as co-regulation groups. The adaptive landscape is reduced to $p + 1$ dimensions – the number of co-regulation groups plus the flux. As in case of regulation between all enzymes, when there is only positively co-regulated or independent enzymes, the flux can increase indefinitely, whereas flux has a maximum if there is at least two negatively co-regulated enzymes (figures 2, 3, 4 and S1).

(vi) When there is co-regulation groups with competition between all enzymes, the constant total concentration complicates the relationship between enzyme concentrations. From the previous cases, we know that the number of degrees of freedom is between 1 and $n-1$. We might hypothesize that the number of df is $p-1$ – the number of co-regulation groups minus the relationship between enzyme concentrations due to constant total concentration –, but this is not coherent with the extreme case of co-regulation between all enzyme (indeed $p=1$ leading to zero df). Another hypothesis for the number of df is to consider each $E^q$ (minus one due to the constant total concentration) and each $\tau_q$ separately, leading to formula $p + \sum_{q=1} \mathbf{1}_{\{\theta_q \neq 0\}} - 1$. This formula is coherent with extreme cases, but do not take account of the complex relationship between sums of enzyme concentrations of a group $E^q$ and the driving variables $\tau_q$ – the same relationship that prevents us to find the effective equilibrium. This case is not very intuitive, because it occurs in a multidimensional space and because the relationship between enzyme concentrations depends on three different factors – co-regulation within groups, independence between groups, competition between all. Note that if a co-regulation group is initially as its theoretical equilibrium (equation S45), the system loses one df as previously. Whatever the degrees of freedom, the adaptive landscape is a projection in lower dimensions of the dome of competition, and its shape is also a dome (figures 5 and 6). This projection depends on the number of co-regulation groups, and each new co-regulation between two previously independent groups diminish the dimension number of the projection.

The equilibrium distribution in case of competition with co-regulation groups depends on the kind and number of co-regulation groups. When all enzymes are co-regulated, the distribution seems random, but actually depends on two points, initial concentrations $\mathbf{E^0}$ and theoretical equilibrium $1/\mathbf{B}$, that fixed the trajectories in the $n$-dimensional space of enzyme concentrations (figures 7B and S2). Each effective equilibrium is the maximum of the projection of the global of competition in this specific condition. With less restriction, *i.e.* with more co-regulation groups, the concentration trajectories can move more freely in the space. Thus, the ensemble of effective equilibrium for different initial concentrations exhibits a more visible pattern (figures 7D, S3, 7F S5, 7H and S4).

Thus the adaptive landscape arises in lower dimensions in the multidimensional space $(\mathbf{E}, J)$, depending on the degrees of freedom that give the different constraints. Its shape is determined by the pseudo-activities, but also the co-regulation coefficients and the initial concentrations, according to the applied constraint.
